# First-in-class [^18^F]-PET radiotracers for GPR39: A platform for noninvasive target engagement imaging *in vivo*

**DOI:** 10.64898/2026.11.06.764940

**Authors:** Bhuvanachandra Bhoopal, Ryan W. Fitzgerald, Krishna K. Gollapelli, Mack Miller, Naresh Damuka, Chrystal B. Baity, Pritam Roy, Paul J. Myburgh, Dongqin Zhu, Pooja Jadiya, Rita Cervera-Juanes, Kiran K. Solingapuram Sai

## Abstract

Disrupted zinc homeostasis is a hallmark of Alzheimer’s disease. As a key zinc-sensing receptor, GPR39, a G-protein coupled receptor is implicated in neurodegeneration, vascular regulation, and neuropsychiatric disorders, represents an emerging target for molecular imaging. However, the lack of suitable positron emission tomography (PET) radiotracers has limited GPR39 *in vivo* characterization. Quantitative assessment of GPR39 levels *in vivo* helps in understanding its role in physiology and pathology. This will facilitate the development of targeted therapeutics and enable real-time monitoring of treatment response in clinical settings. This study introduces [^18^F]GPR39-2.1 as a PET radiotracer specificity towards GPR39. [^18^F]GPR39-2.1 exhibited excellent serum stability with negligible radio-metabolite formation. *In vitro* cell uptake studies showed reduced tracer uptake (55–70%) upon pharmacological blockade. PET imaging demonstrated rapid brain uptake and efficient with blood–brain barrier penetration, followed by gradual washout. Blockade studies in rats revealed significant reductions in brain uptake (up to ∼90%, \*\*\*\**p*<0.0001), consistent across multiple brain regions. Receptor specificity was further validated in GPR39 knockout mice, which exhibited reduced brain uptake of 76% (\*\**p*<0.002) and 90% (\*\*\**p*<0.0002) in both PET imaging and biodistribution studies, respectively. Moreover, significantly lower brain uptake observed in 5xFAD transgenic mice (PET imaging: 40%, \*\*\**p*<0.0002 and biodistribution: 77%, \*\**p*<0.002) indicates sensitivity of [^18^F]GPR39-2.1 to disease-associated changes of GPR39. Biodistribution analysis revealed favorable pharmacokinetics. Autoradiography studies further confirmed the specificity of [^18^F]GPR39-2.1 towards GPR39, corroborating PET imaging and biodistribution studies. These findings establish [^18^F]GPR39-2.1 as a highly specific, stable, and brain-penetrant PET radiotracer for imaging GPR39 *in vivo*. This work supports the utility of [^18^F]GPR39-2.1 as a noninvasive molecular imaging agent for neurological and systemic diseases associated with GPR39 dysregulation. Overall, this work establishes a translational framework for investigating GPR39 biology and facilitates its clinical translation as a diagnostic and therapeutic target through [^18^F]GPR39-2.1.

## 1. Introduction

Zinc is an essential trace metal that supports a wide range of biological functions, acting as a signaling molecule, catalytic cofactor, and structural element in the human body. It regulates numerous cellular processes and is needed for several enzymatic reactions [1–4]. Zinc (Zn^2+^) deficiency, which affects millions of people worldwide, has been linked to impaired immunity, developmental abnormalities, inflammatory conditions, and neurological dysfunction [5–7]. In the brain, zinc is highly concentrated within synaptic vesicles of glutamatergic neurons, where it helps regulate neurotransmission and contributes to normal neural function [8–10]. While physiological levels of zinc are neuroprotective, excessive accumulation can become toxic, leading to neuronal damage and disrupted brain function [11–14]. Consequently, disturbances in zinc homeostasis have been associated with several neurological and neurodegenerative disorders, including depression, epilepsy, cerebral ischemia, Alzheimer’s disease (AD), Parkinson’s disease, amyotrophic lateral sclerosis (ALS), and schizophrenia [15–18].

In AD and related dementias, zinc plays a key role in neurodegenerative processes such as tau hyperphosphorylation, neuroinflammation, and oxidative stress [19–21]. The zinc-sensing G protein coupled receptor 39 (GPR39) plays a central role in maintaining zinc homeostasis, and its dysregulation has been directly linked to amyloid-β (Aβ) accumulation and tau pathology [19, 22]. Zinc dyshomeostasis is a well-established driver of AD pathology, and the zinc-sensing GPR39 sits at the center of this disrupted regulatory network [20, 23]. Under physiological conditions, zinc activates GPR39 to support synaptic integrity, neuronal survival pathways, and homeostatic control of excitatory signaling [22, 24]. In AD, Aβ sequesters zinc into neurotoxic oligomeric complexes, depleting synaptic zinc and impairing GPR39-dependent signaling [25, 26]. Loss of GPR39 contributes to synaptic dysfunction, heightened vulnerability to oxidative and inflammatory stress, and progressive neurodegeneration [25, 27]. Restoring GPR39 function either through targeted agonists or by using molecular imaging to guide therapeutic engagement represents a compelling strategy to counteract zinc dyshomeostasis and intervene directly in a convergent pathway underlying AD progression [24].

GPR39 has emerged as an important modulator of intracellular signaling in both the central nervous system and peripheral tissues [22, 28]. GPR39 activation has been implicated in neuronal survival [22], synaptic plasticity [24], and metabolic regulation [29]. There is further evidence linking the receptor to epilepsy [30], depression [19], AD [20], and cardiovascular related disease [31]. Despite increasing interest in GPR39 biology, its regional distribution, regulation, and pharmacology in the living brain remain poorly characterized [22].

Beyond neurodegeneration, GPR39 also plays a role in other diseases and disorders. In alcohol use disorders (AUD), chronic alcohol exposure disrupts zinc homeostasis, particularly in hippocampal and cortical circuits where zinc is co-released with glutamate [32]. Alcohol reduces synaptic zinc availability, alters vesicular zinc transport, and increases oxidative stress. This leads to impaired activation of GPR39, which normally supports synaptic stability, neuroprotection, and homeostatic excitability control [15]. Reduced GPR39 signaling contributes to the hyperexcitability, anxiety-like behavior, and impaired neuroplasticity characteristics of alcohol withdrawal [33]. Preclinical studies show that GPR39 agonists can reverse alcohol-induced deficits in brain-derived neurotrophic factor signaling, normalize stress-related pathways, and reduce withdrawal-associated anxiety, highlighting the importance of GPR39 [33].

GPR39 also plays important role in the vascular dysfunction that underlies hypertension [34, 35]. GPR39 is expressed in vascular endothelial and smooth-muscle tissues, where it regulates inflammatory signaling, vascular tone, and endothelial resilience [34]. In hypertension, particularly angiotensin II driven models, endothelial injury and oxidative stress impair GPR39-dependent homeostatic signaling [35]. Reduced GPR39 activation contributes to heightened vascular inflammation, impaired nitric oxide bioavailability, and increased susceptibility to hypertensive damage [24, 35]. Conversely, restoring GPR39 mediated signaling has been shown to protect against endothelial dysfunction, positioning this pathway as a mechanistically relevant and potentially targetable axis in the pathophysiology of hypertension [24, 35].

Given the importance of GPR39, it is emerging as a key regulator of both vascular and neuronal homeostasis, with growing evidence supporting their involvement in diverse pathological conditions, including AD, hypertension, and AUD. While GPR39 agonists show significant therapeutic potential for cardiovascular disease [35–37], alcohol use disorder [33], neuroprotection [30], and cognitive recovery [30], the field lacks an non-invasive tool to quantify real-time GPR39 levels in pathological conditions or monitor target engagement during treatment.

Positron emission tomography (PET) enables quantitative measurement of receptor availability and pharmacokinetics in living subjects and has played a critical role in advancing the study of GPCR systems [38, 39]. A GPR39 PET tracer would enable noninvasive visualization and quantification of receptor engagement *in vivo*, facilitating translational evaluation of GPR39 signaling in both peripheral and central disease settings beyond that of only neurodegeneration. We recently reported the first successful PET tracer for imaging GPR39, [^11^C]TMN-OMe, a radiolabeled analog of the GPR39 agonist TM-N1234 [40]. However, due to its moderate brain uptake and the short half-life of [^11^C] (20.4 min), constrained kinetic modeling, and requiring an on-site cyclotron [41], motivated the development of [^18^F]-analogs [42]. [^18^F]-tracers offer several advantages over [^11^C]-tracers for PET imaging research, including a longer half-life (109.8 min), higher achievable injected activities, and greater flexibility in imaging protocols, all of which facilitate more robust kinetic analyses and improve suitability for clinical translation [41, 43]. TC-G1008 is a well-characterized high affinity GPR39 ligand (EC_50_ = 0.8 nM in humans) [44, 45]. Our three GPR39 [^18^F]-based tracers were modeled based on the lead compound GTPL7803 (TC-G1008 derivative) [28, 45].

In the present study, we report the design and synthesis of three [^18^F]-based tracers for GPR39, followed by the evaluation of [^18^F]GPR39-2.1 through *in vitro* and *in vivo* studies, including serum stability, cell uptake, PET imaging, tissue biodistribution, and autoradiography. By leveraging the favorable half-life of [^18^F] and a high affinity scaffold, this work aims to identify a lead PET tracer capable of robust, noninvasive GPR39 imaging, thereby advancing translational research into this emerging therapeutic target.

## 2. Results

### 2.1 In silico analysis

Molecular docking analysis was performed to evaluate the binding affinity of the designed ligands against the GPR39. The lead molecule (GTPL7803) and its derivatives (GPR39-2.1, GPR39-2.2, and GPR39-2.3) bind within the same pocket as human GPR39 with minimal deviation in pose **(Figure 1).** Among the tested compounds, GPR39-2.1 exhibited the highest binding affinity with a docking score of −9.94 kcal/mol. The lead molecule demonstrated a slightly lower binding affinity, with a docking score of −9.57 kcal/mol. The GPR39-2.3 compound yielded a docking score of −9.18 kcal/mol. GPR39-2.2 exhibited the lowest binding affinity among the evaluated ligands, with a docking score of −8.46 kcal/mol. Overall, the docking results indicate that all tested ligands occupy the same binding pocket and exhibit comparable binding affinities relative to the lead molecule. The BOILED-Egg model analysis revealed that all derivatives (GPR39-2.1, GPR39-2.2, and GPR39-2.3), along with the lead molecule have a high probability of blood–brain barrier (BBB) permeation with good pharmacokinetics (**Table S1**). This suggests that these compounds possess physicochemical properties favorable for central nervous system (CNS) accessibility. Structural modifications introduced in the lead molecule did not negatively impact their ability to cross the BBB. Instead, these derivatives retained optimal balance between lipophilicity and polarity, which is critical for passive diffusion across the BBB. Overall, the BOILED-Egg model predictions demonstrate that all evaluated ligands exhibit favorable BBB penetration profiles, reinforcing their potential suitability for targeting GPR39 in CNS-related applications *in vivo*.

**Figure 1.**
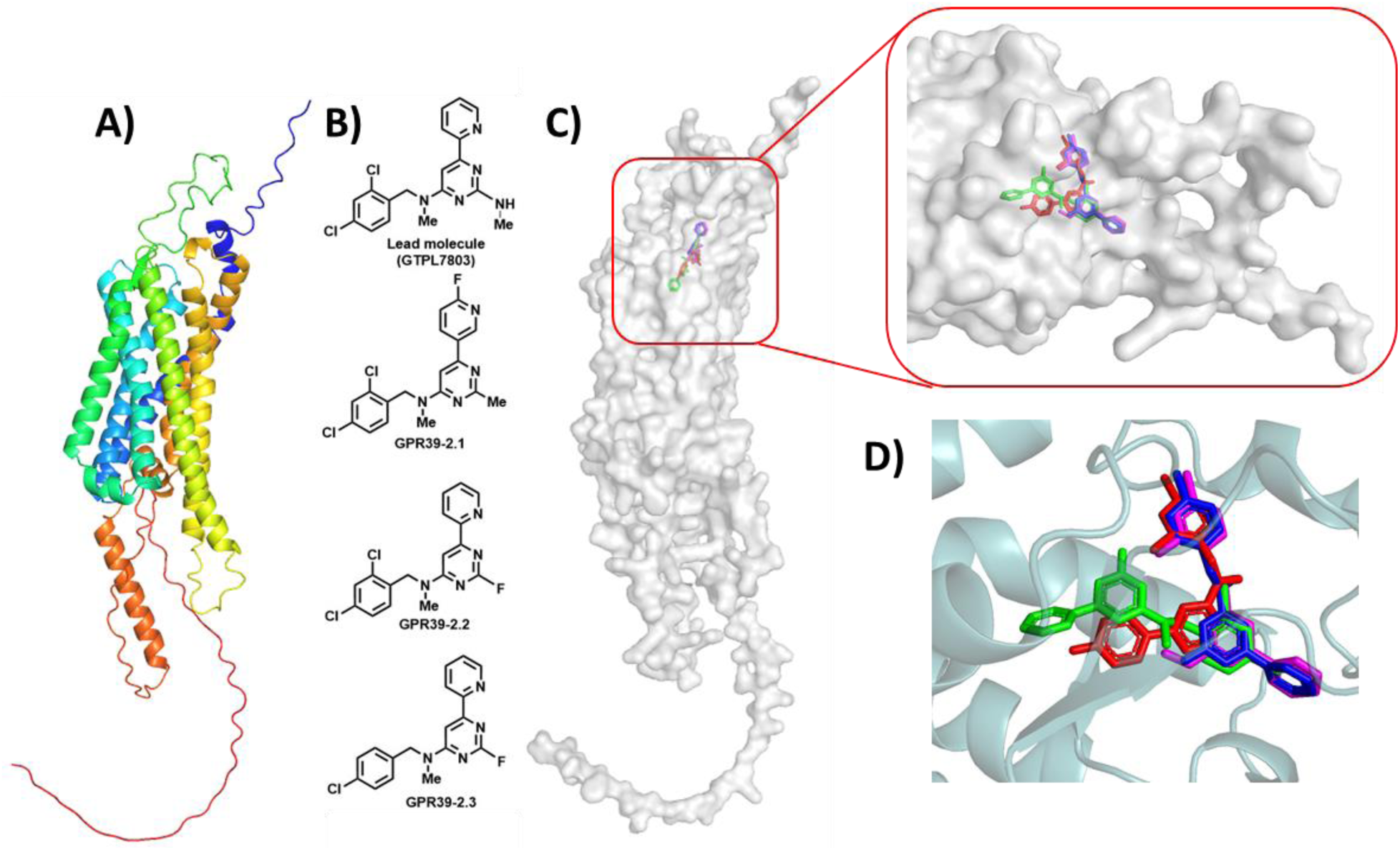
Molecular docking studies of GPR39. **A)** 3D structure of GPR39 protein in cartoon representation. **B)** Structures of small molecules used for molecular docking studies. **C)** GPR39 protein in surface representation with docked ligands in the binding pocket. **D)** Lead molecule (GTPL7803; magenta) and its derivatives GPR39-2.1 (red), GPR39-2.2 (green), and GPR39-2.3 (blue) docked to GPR39 (cartoon representation).

### 2.2 Synthesis of GPR39-2.1 (6), GPR39-2.2 (11), and GPR39-2.3 (16)

[^18^F]GPR39 precursors and standards were synthesized as shown in **Scheme 1**. Precursors **5**, **10**, and **15** were synthesized with 42%, 56%, and 42% chemical yields, respectively as off-white solids. The nonradioactive fluorine-19 standards **6**, **11**, and **16**, were synthesized with 67%, 58%, and 66% chemical yields, respectively furnished as off-white solids (compounds chemical synthesis is given as supplemental methods and characterization data is given as **Fig. S1-S21**)

**Scheme 1.**
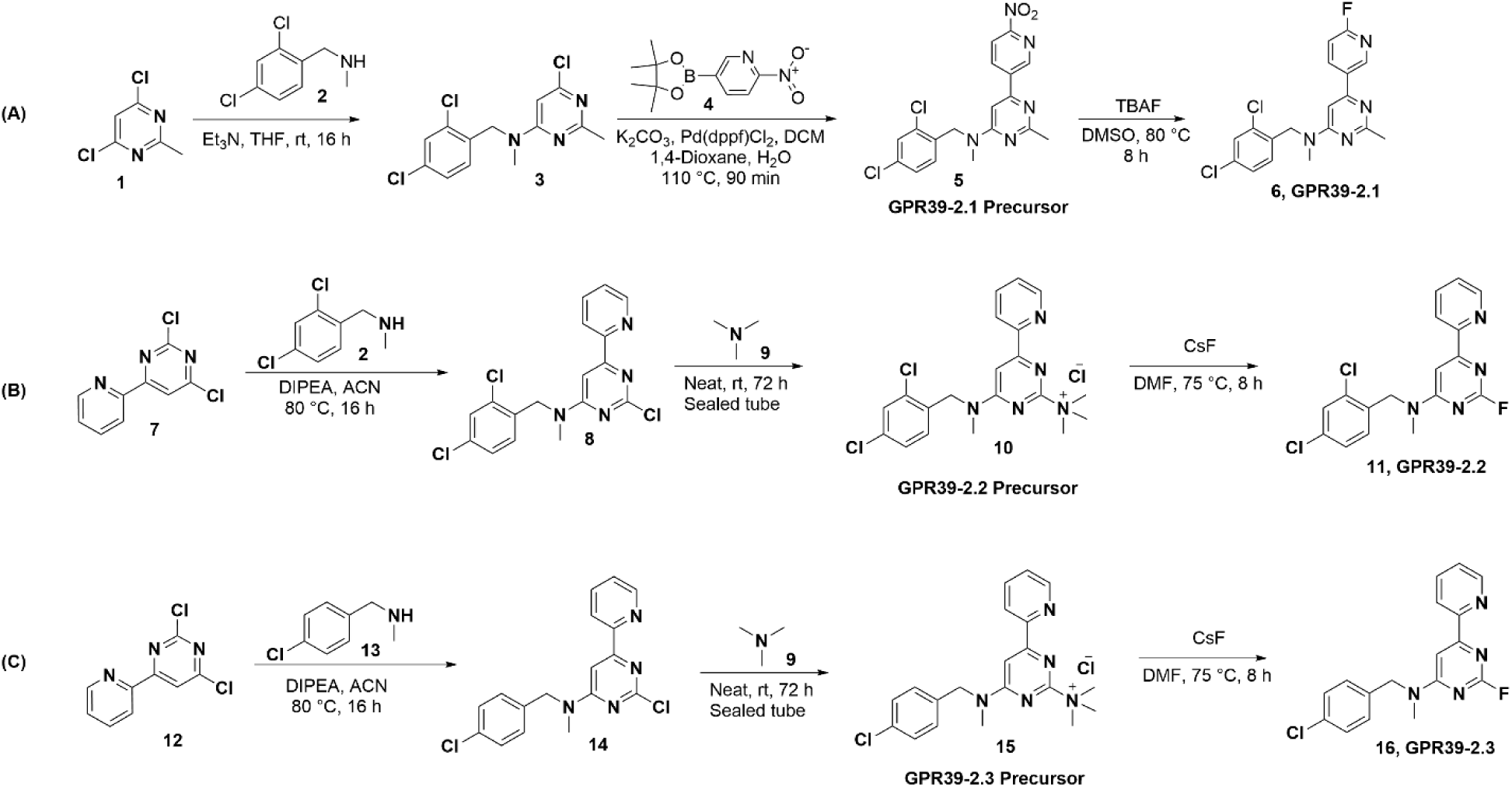
(A) Synthesis of GPR39-2.1 precursor (**5**) and GPR39-2.1 (**6**); (B) Synthesis of GPR39-2.2 (**10**) precursor and GPR39-2.2 (**11**); (C) Synthesis of GPR39-2.3 precursor (**15**) and GPR39-2.3 (**16**).

### 2.3 Radiochemistry

The complete radiochemical synthesis (**Scheme 2**), including labeling, semi-preparative HPLC purification, and final formulation was accomplished within 90 min for [^18^F]GPR39-2.1 ([^18^F]**6**), and 60 min for [^18^F]GPR39-2.2 ([^18^F]**11**) and [^18^F]GPR39-2.3 ([^18^F]**16**) (**Fig. S22-S24**). A molar activity of 242.0 ± 56.6 GBq/µmol and radiochemical yield of 15 ± 5% (n > 15) was obtained for [^18^F]GPR39-2.1. Molar activity of [^18^F]GPR39-2.2 was 23.7 ± 9.0 GBq/µmol with a radiochemical yield of 6.1 ± 0.5% (n > 4), while [^18^F]GPR39-2.3 had a molar activity of 18.2 ± 5.0 GBq/µmol with a radiochemical yield of 10.0 ± 0.5% (n > 4). All reported molar activity values were decay corrected. The radiochemical purity for all three compounds were > 99%, assessed by total peak area on the analytical HPLC and radiochemical identity was confirmed via co-injection with nonradioactive standards corresponding to each radiotracer (**Fig. S25-S27**). The high radiochemical purity and consistent HPLC retention times confirm the reliability and reproducibility of the labeling process.

**Scheme 2.**
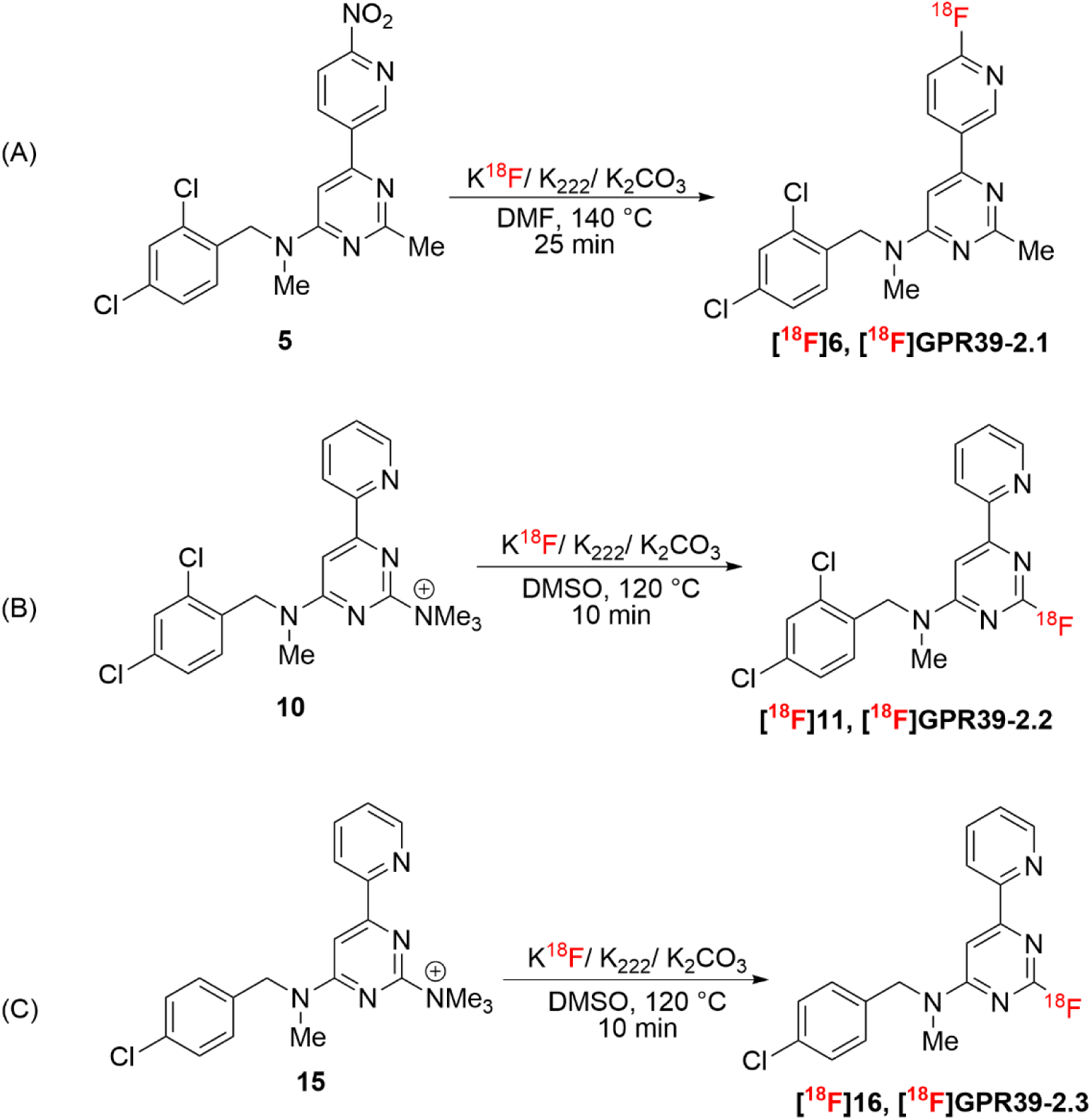
Radiochemical synthesis of **(A)** [^18^F]GPR39-2.1, **(B)** [^18^F]GPR39-2.2, and **(C)** [^18^F]GPR39-2.3.

### 2.4 Serum stability

Stability of radiotracers in human serum was evaluated on analytical HPLC equipped with radio detector. All three radiotracers exhibited high stability, with the parent compound remaining at all time points. Radiochemical purity (RCP) exceeded 99% throughout the incubation period, and no additional radioactive peaks were observed in the chromatograms (**Figure 2**). [^18^F]GPR39-2.2 (**11**) and [^18^F]GPR39-2.3 (**16**) exhibited lower molar activities compared to [^18^F]GPR39-2.1 (**6**), ultimately prioritizing [^18^F]GPR39-2.1 for further *in vitro* and *in vivo* studies over the other analogs.

**Figure 2.**
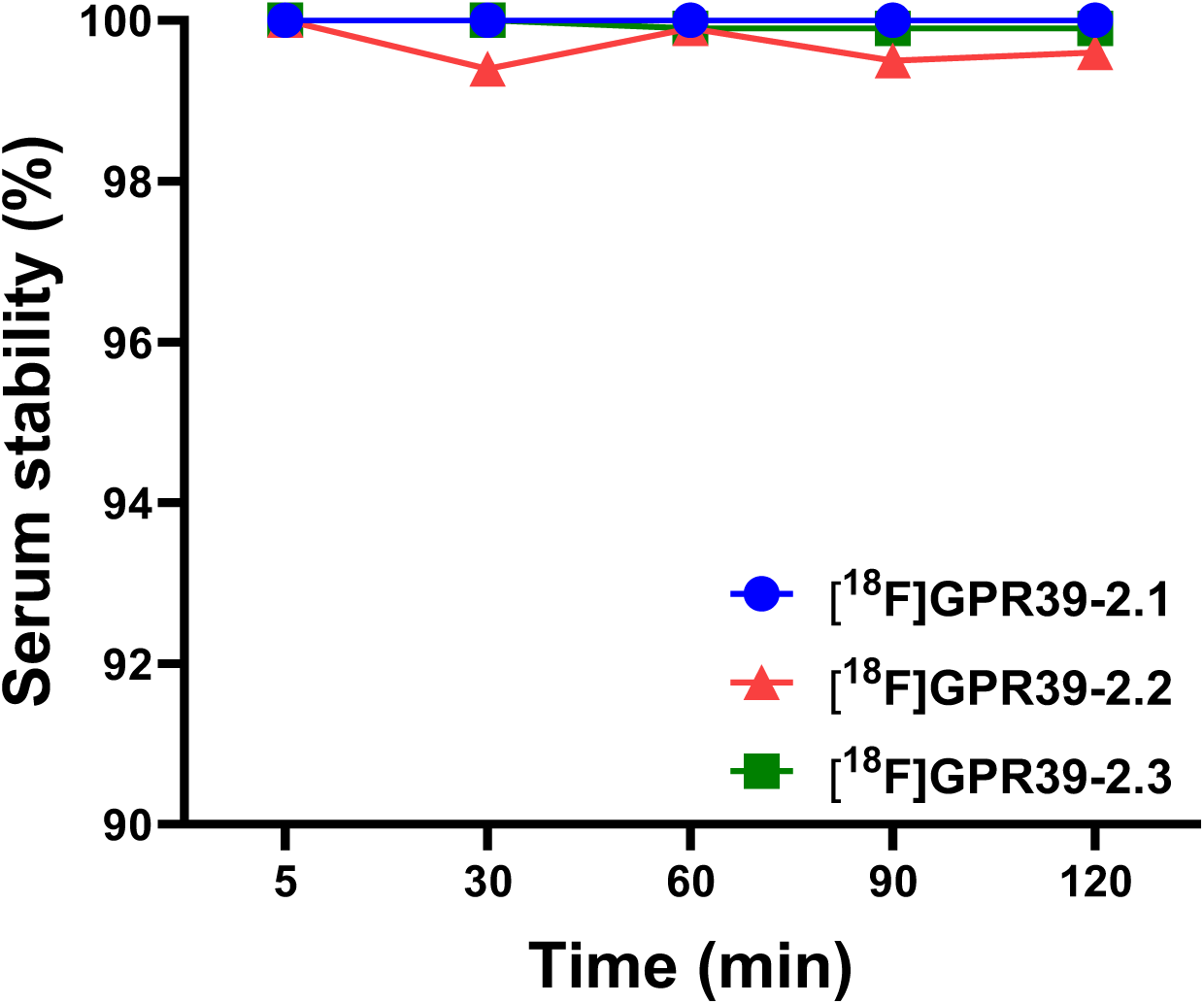
Serum stability of [^18^F]GPR39-2.1, [^18^F]GPR39-2.2, and [^18^F]GPR39-2.3.

### 2.5 Inositol Phosphate 1 (IP1) assay

To evaluate the effect of GPR39-2.1 on the activation of GPR39, a concentration–dependent activation assay measuring IP1 levels was performed across a range of ligand concentrations (0.5 nM to 1 mM). The response was quantified as a homogeneous time-resolved fluorescence (HTRF) ratio and plotted against the compound concentration (**Figure 3**). At low concentrations (>500 nM), GPR39-2.1 elicited minimal to no effect, suggesting that the system is insensitive to ligand at sub-threshold concentrations. Beginning around 5 µM, a clear concentration-dependent activation becomes evident and the sharp decrease in HTRF ratio was observed until 1 mM, indicating a transition into the dynamic range of the assay. This steep portion of the curve suggests that GPR39-2.1 exerts a modulatory effect within this concentration window. The data exhibit a sigmoidal dose–response relationship characteristic of pharmacological activation of GPR39. The midpoint of the response lies between 5 µM and 50 µM, with an estimated EC_50_ of 20.36 µM. Overall, these results demonstrate that GPR39-2.1 activates GPR39 in a dose-dependent manner.

**Figure 3.**
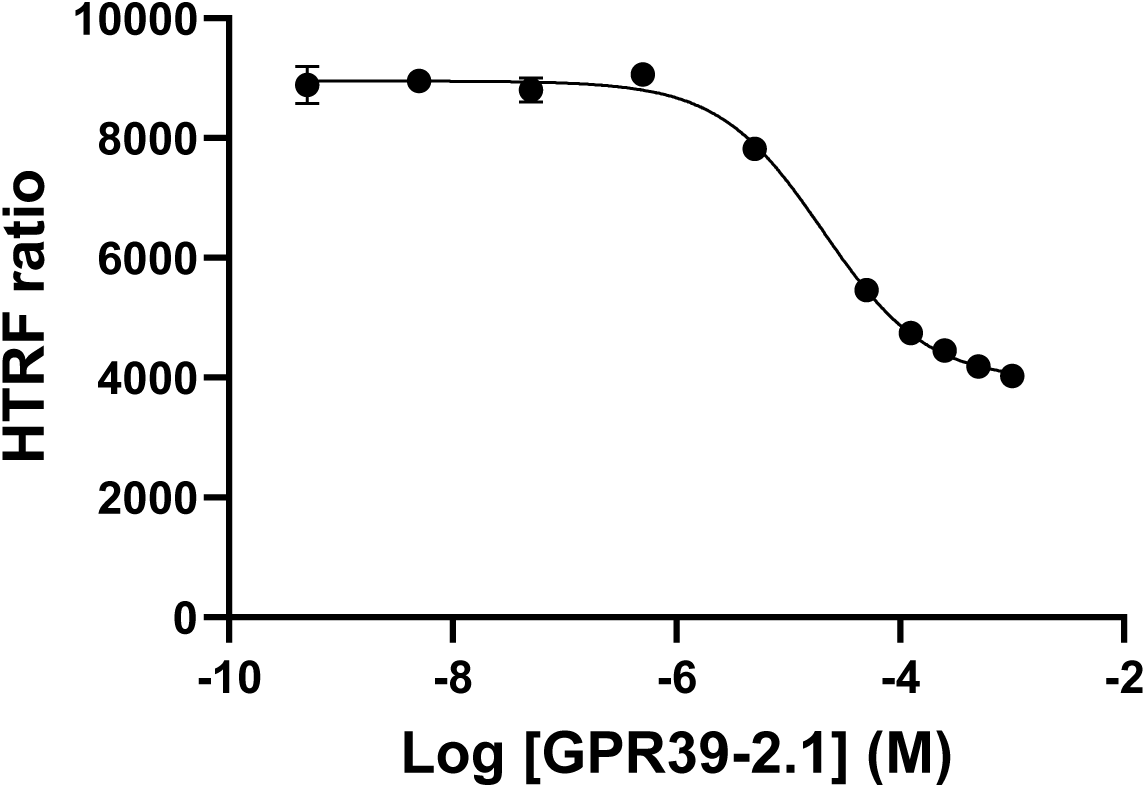
GPR39 activation measured by IP1 accumulation. MC38 cells expressing human GPR39 were incubated with GPR39-2.1 (0.5 nM -1 mM) for 2 h at 37 °C. Data were fit by nonlinear regression to determine EC_50_ value in GraphPad Prism v10.3.1. Data represent mean ± SD of independent experiments performed in triplicate.

### 2.6 GPR39 cell uptake study

The cellular uptake of [^18^F]GPR39-2.1 was evaluated in BT-20 (human breast cancer), MC38 (murine colon cancer), PC3 (human prostate cancer), and U87-MG (human glioblastoma) cell lines to determine intracellular accumulation of the radiotracer and its efficiency in crossing cellular membranes. As shown in **Figure 4A**, the radiotracer exhibited time-dependent uptake across all tested cell lines, with maximum accumulation observed from 60 min onwards in PC3 and U87-MG cells. To assess the specificity of [^18^F]GPR39-2.1 toward GPR39, blocking studies were performed using TC-G1008 and TM-N1324, which are established GPR39 agonists. Pre-incubation of cells with these agonists led to a marked reduction in radiotracer uptake, indicating successful competition for receptor binding sites. Notably, all cell lines demonstrated a significant decrease in uptake under blockade conditions (**Figure 4B**). Quantitatively, the presence of blocking agents resulted in an approximate 55–70% reduction in tracer uptake (*p* ≤ 0.03) across all tested cell lines, confirming receptor-mediated internalization. This substantial decrease highlights the specificity of [^18^F]GPR39-2.1 for GPR39 and minimizes the likelihood of non-specific uptake. Collectively, the *in vitro* cellular uptake studies conducted in both human and murine cancer cell lines including glioblastoma, breast, colon, and prostate models demonstrate that [^18^F]GPR39-2.1 exhibits specific and receptor-mediated uptake at the cellular level.

**Figure 4.**
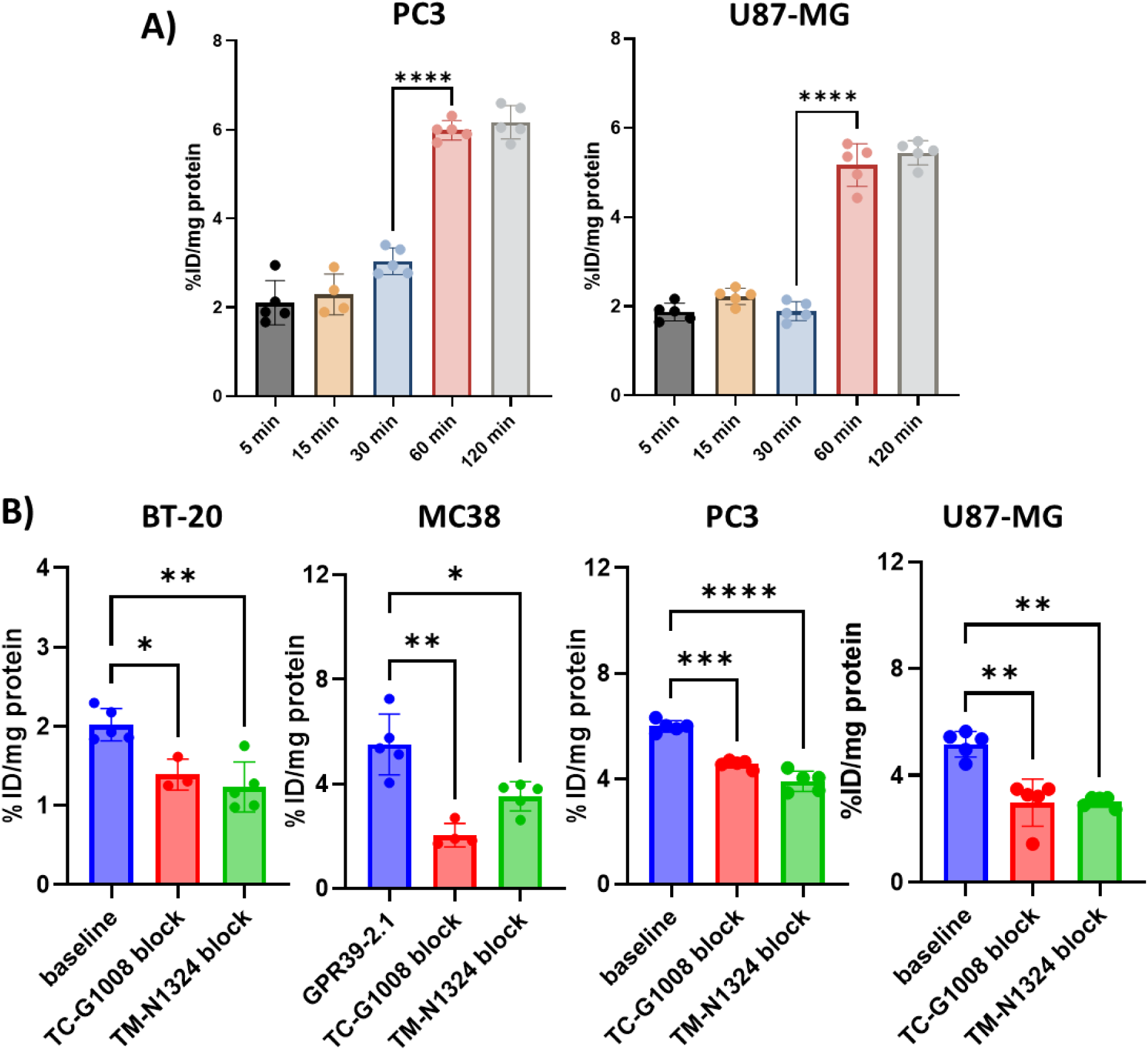
*In vitro* cell uptake assay of [^18^F]GPR39-2.1. **A)** Time-dependent cell uptake kinetics of [^18^F]GPR39-2.1 in PC3 and U87-MG cell lines (n=5/time point). **B)** *In vitro* cell uptake assays of [^18^F]GPR39-2.1 in BT-20, MC38, PC3 and U87-MG cell lines at 60 min under baseline and blockade conditions (TC-G1008 and TM-N1324) (n=5/group). Statistical analysis was performed using two-way ANOVA, \*\*\*\**p*<0.0001, \*\**p*<0.002, \**p*<0.03

### 2.7 In vivo microPET/CT imaging

*In vivo* PET/CT imaging was performed to evaluate brain uptake, and receptor specificity of [^18^F]GPR39-2.1 (**Figure 5**). Whole body image showed favorable kinetics over 120 min (**Figure S28A**). Time activity curves (TACs) showed rapid brain uptake within 5 min, ∼50% of maximal activity at 30 min, and a gradual washout over 120 min (**Figure S28**). Dynamic PET imaging in BALB/c revealed brain uptake of the radiotracer, indicating efficient BBB penetration. Quantitative analysis of brain uptake showed that pre-treatment with TC-G1008 resulted in a marked reduction in tracer uptake (56.2% decrease at 60 min, \**p* < 0.03) compared to baseline condition, demonstrating receptor-specific binding (**Figure 5A**). PET imaging in rats revealed differential regional brain uptake of the radiotracer under baseline and blockade conditions (**Figure 5B**). Significant decrease (\*\*\*\**p*<0.0001) in uptake under blockade conditions was consistent across the cortex (90.9% decrease), hippocampus (90.5% decrease), and in whole brain (90.5% decrease), supporting selective engagement of GPR39 by [^18^F]GPR39-2.1 (**Figure 5C**).

**Figure 5.**
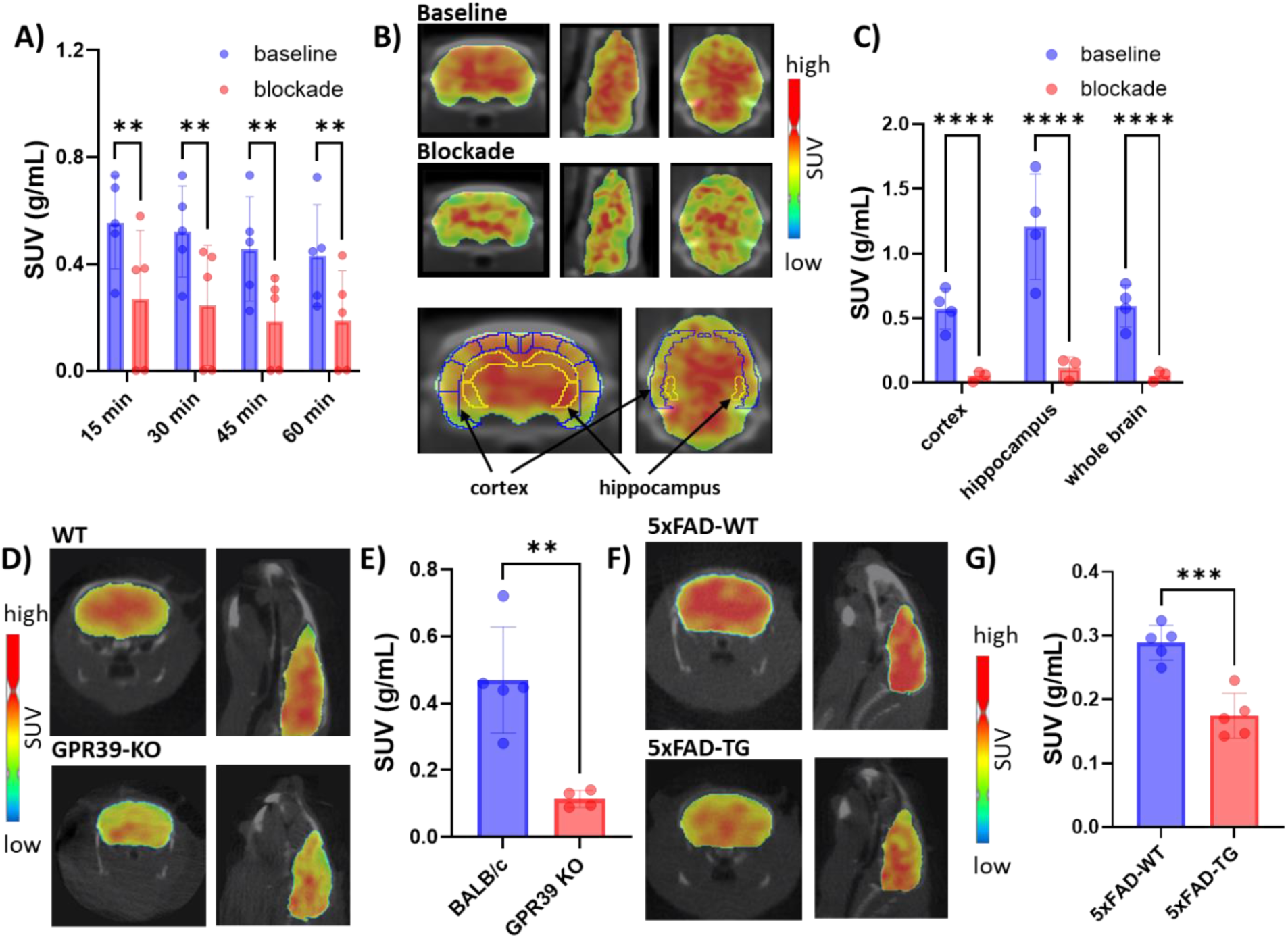
MicroPET/CT imaging of [^18^F]GPR39-2.1. **(A)** Brain SUV quantification from microPET/CT images of [^18^F]GPR39-2.1 in BALB/c mice (n = 5/group) under baseline and blockade (TC-G1008) conditions. Statistical analysis was performed using two-way ANOVA, \*\**p*<0.002. **(B)** Representative microPET/CT images showing regional brain uptake of [^18^F]GPR39-2.1 **(C)** and its SUV quantification in Wistar IGS rats (n = 4/group) under baseline and blockade (TC-G1008) conditions. Statistical analysis was performed using two-way ANOVA, \*\*\*\**p*<0.0001. **(D)** Representative microPET/CT images showing brain uptake of [^18^F]GPR39-2.1 **(E)** and its SUV quantification in wild type mice (n=5) and GPR39-KO mice (n = 4). Statistical analysis was performed using unpaired two-tailed t-test, \*\**p*<0.002. **(F)** Representative microPET/CT images showing brain uptake of [^18^F]GPR39-2.1 **(G)** and its SUV quantification in 5xFAD wild type (WT) and transgenic (TG) mice (n = 5/group). Statistical analysis was performed using unpaired two-tailed t-test, \*\*\**p*<0.0002.

In addition, comparative imaging between wild type and GPR39 knock out (KO) mice revealed a significant decrease in SUV values under blocking conditions compared to baseline (**Figure 5D**). Quantitatively, 75.6% reduced brain uptake was observed in KO mice (\*\**p*<0.002), indicating specific receptor-mediated uptake rather than nonspecific binding (**Figure 5E**). Applicability of the radiotracer in disease models was well supported by the results of PET imaging in AD mouse models (5xFAD). Transgenic 5xFAD mice exhibited reduced brain uptake compared to wild type mouse models (**Figure 5F**). Quantitatively, 39.6% reduced brain uptake was observed in 5xFAD transgenic mice compared to wild type (\*\*\**p*<0.0002) (**Figure 5G**), suggesting reduced GPR39 levels in transgenic mice.

The PET imaging data obtained from higher rodent models (rats) showed consistent brain uptake patterns and effective reduction upon receptor blockade, confirming relevance across species. Similarly, comparative studies demonstrated reduced tracer uptake in KO models lacking GPR39 expression, providing additional validation of target specificity. Reduced tracer uptake in 5xFAD transgenic mice supports in utility in disease context. Overall, PET imaging results demonstrate that [^18^F]GPR39-2.1 exhibits favorable *in vivo* characteristics, including efficient brain penetration, robust and regionally consistent uptake, and significant reduction under blocking conditions. These findings strongly support the specific and receptor-mediated binding of the radiotracer to GPR39, highlighting its potential for noninvasive imaging of GPR39 *in vivo*.

### 2.8 Tissue biodistribution studies

The tissue biodistribution profile of [^18^F]GPR39-2.1 was evaluated at different time points (5-120 min post-injection) across major organs to assess tissue uptake, clearance kinetics, and overall pharmacokinetic behavior (**Table 1**). The tracer demonstrated rapid systemic distribution, with moderate initial uptake in blood (3.77 ± 0.38 %ID/g at 5 min) that slightly increased at early time points before gradually decreasing to 1.31 ± 0.28 %ID/g at 120 min, indicating progressive clearance from circulation. A similar trend was observed in the brain, with uptake increasing from 2.34 ± 0.25 %ID/g at 5 min to a peak of 3.56 ± 0.42 %ID/g at 60 min, followed by a decline at later time points, suggesting transient blood–brain barrier penetration. Among all organs, the kidney exhibited the highest uptake, reaching 17.00 ± 2.35 %ID/g at 60 min, followed by a gradual decrease to 7.95 ± 2.13 %ID/g at 120 min, indicating renal clearance as the primary excretion pathway. The liver also showed relatively high uptake, peaking at 6.80 ± 0.63 %ID/g at 60 min, which declined to 2.73 ± 0.36 %ID/g at 120 min, suggesting partial hepatobiliary involvement in tracer clearance. The lungs displayed moderate early uptake (3.63 ± 0.73 %ID/g at 5 min) with rapid washout over time, decreasing to 0.91 ± 0.62 %ID/g at 120 min. Similarly, heart uptake increased initially to 3.24 ± 0.50 %ID/g at 60 min, followed by a decline, reflecting transient perfusion-related distribution. Low to moderate uptake was observed in spleen, pancreas, and muscle, with values remaining relatively stable or gradually decreasing over time, indicating minimal nonspecific retention. Bone uptake remained consistently low throughout the study (≤0.65 %ID/g), suggesting minimal defluorination and good radiotracer stability. Overall, the biodistribution data indicates that the radiotracer exhibits favorable pharmacokinetics characterized by efficient systemic distribution, moderate brain penetration, predominant renal clearance, and low nonspecific accumulation in non-target tissues.

**Table 1.** Biodistribution of [^18^F]GPR39-2.1 in BALB/c mice (n = 5/time point) *^a^*.

| **Organ** | **5 min** | **30 min** | **60 min** | **90 min** | **120 min** |
|---|---|---|---|---|---|
| Blood | 3.77 ± 0.38 | 4.56 ± 0.41 | 4.82 ± 0.58 | 3.68 ± 0.75 | 1.31 ± 0.28 |
| Brain | 2.34 ± 0.25 | 3.06 ± 0.32 | 3.56 ± 0.42 | 2.59 ± 0.31 | 0.97 ± 0.40 |
| Heart | 1.88 ± 0.43 | 2.91 ± 0.77 | 3.24 ± 0.50 | 1.96 ± 0.77 | 1.08 ± 0.52 |
| Lungs | 3.63 ± 0.73 | 3.23 ± 1.27 | 2.30 ± 0.28 | 0.92 ± 0.43 | 0.91 ± 0.62 |
| Liver | 6.26 ± 0.80 | 6.31 ± 0.88 | 6.80 ± 0.63 | 3.79 ± 0.99 | 2.73 ± 0.36 |
| Spleen | 1.72 ± 0.34 | 1.80 ± 0.38 | 1.90 ± 0.94 | 1.27 ± 0.76 | 1.56 ± 0.60 |
| Kidney | 13.14 ± 1.36 | 15.09 ± 0.90 | 17.00 ± 2.35 | 11.78 ± 1.04 | 7.95 ± 2.13 |
| Pancreas | 1.82 ± 0.48 | 2.28 ± 0.38 | 2.21 ± 0.69 | 1.48 ± 0.32 | 0.82 ± 0.28 |
| Muscle | 0.55 ± 0.29 | 1.03 ± 0.57 | 1.27 ± 0.66 | 0.93 ± 0.38 | 0.83 ± 0.27 |
| Bone | 0.22 ± 0.07 | 0.55 ± 0.39 | 0.61 ± 0.25 | 0.16 ± 0.03 | 0.65 ± 0.35 |

<sup>*a*</sup>Values were reported as percentages of the injected dose per gram of tissue (%ID/g) ± SD.

The tissue biodistribution of [^18^F]GPR39-2.1 was further evaluated under baseline and blocking conditions in Wistar IGS rats, as well as in KO mice and 5xFAD mice, to assess tracer specificity and biological relevance (**Figure 6**). In rats, baseline biodistribution demonstrated moderate-to-high uptake of [^18^F]GPR39-2.1 across several organs. Pre-treatment with TC-G1008 resulted in a significant reduction in tracer uptake across multiple tissues (**Figure 6A**), indicating receptor-mediated binding. Quantitative analysis revealed statistically significant decreases in uptake under blocking conditions in brain (56.5% decrease), supporting the specificity of [^18^F]GPR39-2.1 for GPR39. A consistent trend of reduced uptake was observed across most organs, further validating target engagement *in vivo*. To confirm target specificity at the genetic level, tracer uptake was compared between wild-type and GPR39 KO mice (**Figure 6B**). A clear reduction in brain uptake (89.5%, \*\*\**p*<0.0002) was observed in GPR39 KO animals relative to wild-type controls, indicating that the observed signal is largely dependent on GPR39 levels. Comparative analysis revealed significantly reduced brain uptake (76.8%, \*\**p*<0.002) of the radiotracer in the 5xFAD transgenic mice compared to wild type (**Figure 6C**). These findings suggest that GPR39 levels or accessibility may be modulated in the disease conditions, highlighting the tracer’s potential utility for investigating neurodegenerative pathology. Overall, the tissue biodistribution results demonstrated that [^18^F]GPR39-2.1 exhibits specific, receptor-mediated uptake, as evidenced by pharmacological blockade, genetic knockout validation, and differential uptake in a disease model. These findings collectively support PET imaging studies in both physiological and pathological conditions.

**Figure 6.**
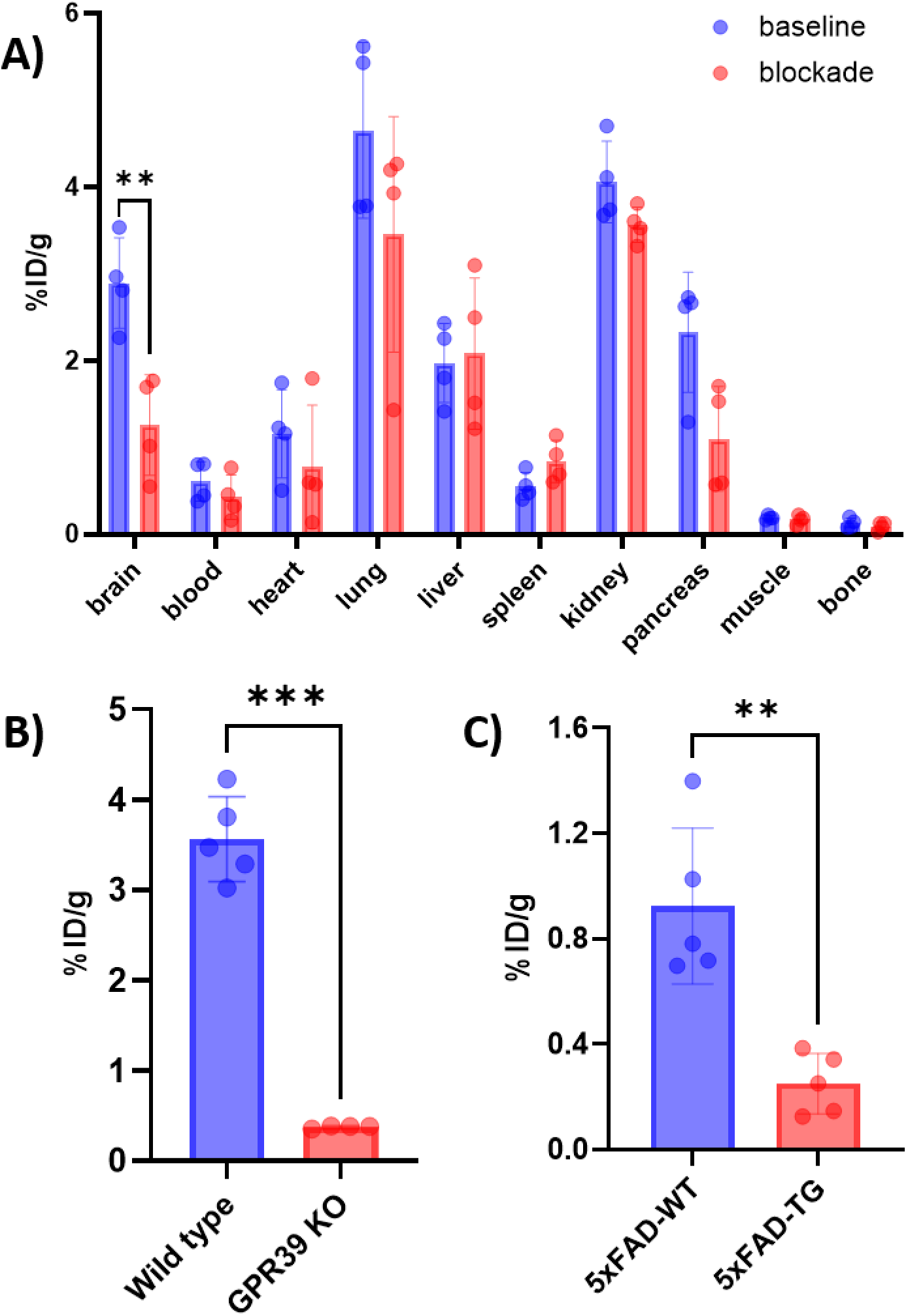
Biodistribution studies of [^18^F]GPR39-2.1. **(A)** [^18^F]GPR39-2.1 uptake in Wistar IGS rats at baseline and blockade (TC-G1008) conditions (n = 4/group) in different organs. Statistical analysis was performed using two-way ANOVA, \*\**p*<0.002. **(B)** [^18^F]GPR39-2.1 brain uptake between wild type (n = 5) and GPR39 KO mice (n = 4). Statistical analysis used unpaired two-tailed t-test, \*\*\**p*<0.0002. **(C)** [^18^F]GPR39-2.1 brain uptake between wild type and transgenic mice of 5xFAD (n = 5/group). Statistical analysis was performed using unpaired two-tailed t-test, \*\**p*<0.002.

### 2.9 Autoradiography

*In vitro* autoradiography was performed to further confirm its specificity toward GPR39. Autoradiography studies demonstrated a reduced radiotracer binding under blockade conditions (TM-N1324 and TC-G1008) compared to baseline, indicating effective competition for receptor binding sites and confirming the specificity of [^18^F]GPR39-2.1 towards GPR39 (**Figure 7A**). Quantitative analysis further supported these observations, with baseline samples showing the highest tracer binding, while pre-treatment with blocking agents resulted in a significant decrease in radiotracer binding (TM-N1324: 62.4% decrease, \*\*\**p*<0.0002; TC-G1008: 79.4% decrease, \*\*\**p*<0.0002) (**Figure 7B**). Overall, these findings demonstrated that [^18^F]GPR39-2.1 exhibits specific and receptor-mediated binding *in vitro*, as evidenced by the significant reduction in signal following receptor blockade. The strong baseline signal combined with effective displacement under blocking conditions highlights the tracer’s specificity for GPR39, corroborating *in vivo* PET imaging and biodistribution studies.

**Figure 7.**
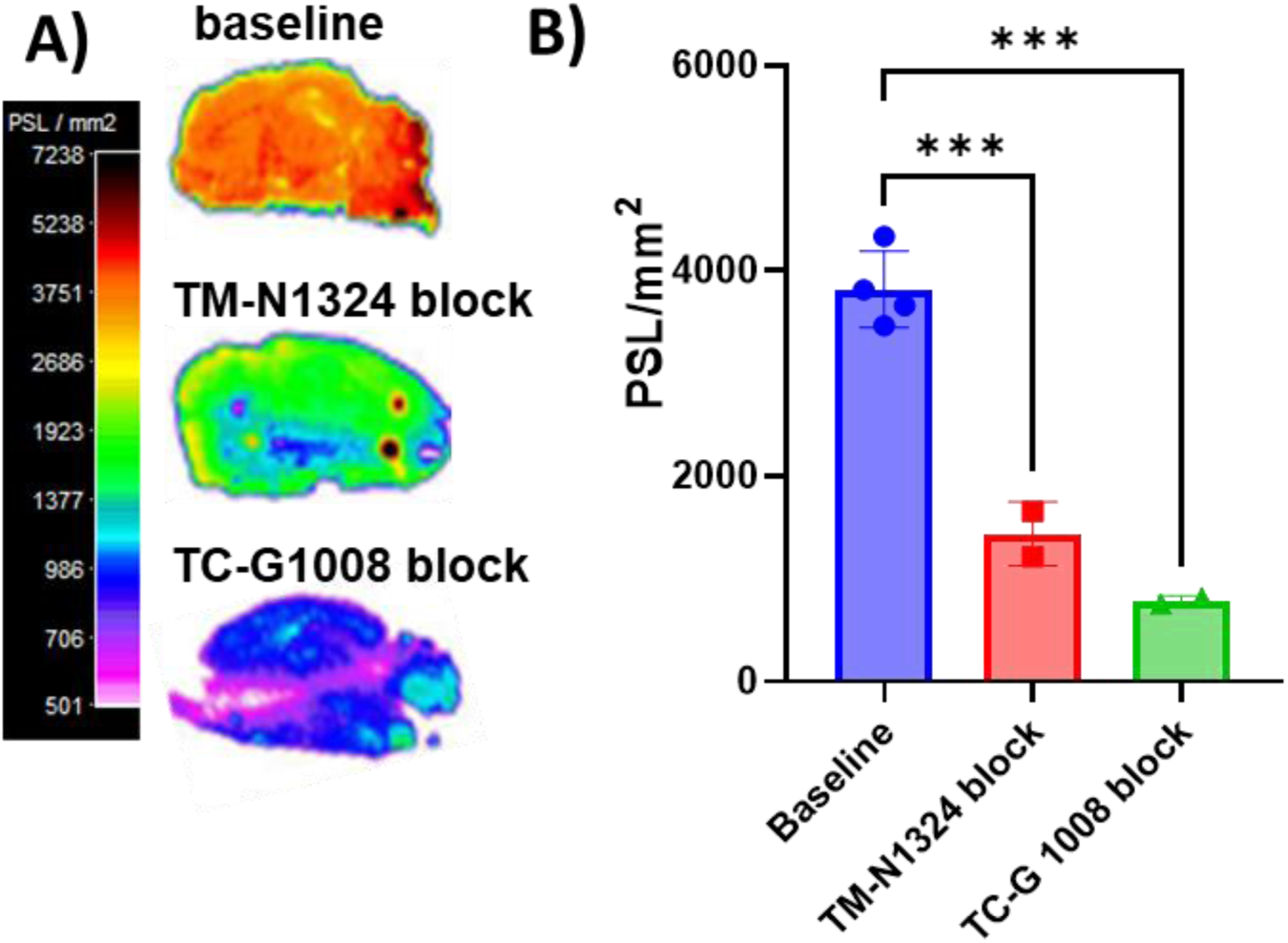
*In vitro* autoradiography studies of [^18^F]GPR39-2.1. (A) Representative *in vitro* autoradiograms and (B) their quantification of [^18^F]GPR39-2.1 binding to BALB/c mice brain tissue sections (n = 4) at baseline and blockade conditions (TM-N1324 or TC-G1008). Statistical analysis was performed using two-way ANOVA analysis, \*\*\**p*<0.0002.

## 3. Discussion

The development of a reliable GPR39-targeted radiotracer is essential to understand the role of GPR39 in of zinc homeostasis and GPR39 involvement in pathology of cardiovascular, neurological, and alcohol use disorders. While our previous work with [^11^C]TMN-OMe provided proof-of-concept for GPR39 imaging, the short half-life of [^11^C] (t½ = 20.4 min) limits its use to facilities with an on-site cyclotron. In this study, we successfully developed and evaluated a series of [^18^F]-labeled radiotracers for the first time and fully characterized them, among which [^18^F]GPR39-2.1 emerged as a highly promising candidate for the noninvasive quantification of GPR39 expression.

The present study integrates *in silico* study, radiochemistry, and comprehensive biological validation to verify [^18^F]GPR39-2.1 as a robust and specific imaging tool to image GPR39 *in vivo*. The transition to [^18^F] (t½ = 109.8 min) was achieved by identifying GTPL7803 as a lead scaffold. Computational docking demonstrated that all designed ligands occupy a conserved binding pocket within GPR39, with GPR39-2.1 exhibiting the highest binding affinity, consistent with its preserved interaction geometry relative to the lead molecule. Importantly, BOILED-Egg predictions indicated that structural refinements did not compromise physicochemical properties critical for BBB permeability, suggesting that rational ligand optimization successfully balanced lipophilicity and polarity. [^18^F]GPR39-2.1 displayed superior molar activity compared to [^18^F]GPR39-2.2 and [^18^F]GPR39-2.3. The high molar activity of [^18^F]GPR39-2.1 is particularly crucial for GPR39 imaging, as GPCRs in the brain often exist in low densities; strong binding with less mass is required for accurate quantification of GPR39. *In silico* data with radiochemistry highlights the BBB penetration capability of three tracers and identifying [^18^F]GPR39-2.1 as the most promising candidate for further evaluation.

The functional and cellular assays further corroborate the *in silico* studies by demonstrating that GPR39-2.1 is a high affinity biologically active modulator of the receptor. The observed sigmoidal dose–response relationship in the IP1 assay confirms receptor activation consistent with pharmacological engagement of GPR39. This functional activity aligns with the receptor-specific uptake observed in multiple human and murine cell lines, where competitive blockade using known agonists resulted in a substantial reduction (55–70%) in radiotracer accumulation. These findings strongly support GPR39 specific cell uptake. Moreover, the excellent serum stability over 120 min and absence of radio-metabolites reinforce the translational robustness of [^18^F]GPR39-2.1. Combination of binding affinity, receptor activation, and selective cellular uptake establishes a coherent structure–activity–function relationship of [^18^F]GPR39-2.1 towards GPR39.

Translational relevance of [^18^F]GPR39-2.1 is validated through *in vivo* PET imaging, biodistribution, and autoradiography studies across multiple models. TACs in BALB/c mice revealed rapid brain uptake of [^18^F]GPR39-2.1 under baseline conditions, indicating efficient penetration across the BBB. Pharmacological blockade with TC-G1008 resulted in a significant reduction (\**p* < 0.03) in tracer uptake (56% at 60 min), confirming receptor-specific binding *in vivo*. GPR39 is widely distributed within the central nervous system, with particularly high expression observed in the hippocampus, amygdala, and frontal cortex [46–49]. Initial metabolite analysis in mice confirms the stability of the tracer. In rats, regional analysis further strengthened these findings, with pronounced reductions >90% (\*\*\*\**p*<0.0001) in the cortex, hippocampus, and whole brain under blockade conditions. This substantial and consistent attenuation across key brain regions highlights the high specificity and selective engagement of GPR39 by the radiotracer *in vivo*. These results demonstrate robust, regionally consistent, and receptor-mediated engagement of GPR39 by [^18^F]GPR39-2.1 across species.

This specificity is further supported by genetic and disease models, where reduced brain uptake in GPR39 knockout mice (76% reduction, \*\**p*<0.002) and 5xFAD transgenic mice (40% reduction, \*\*\**p*<0.0002) directly correlates with altered receptor levels. The decreased uptake observed in the 5xFAD model is consistent with decreased GPR39 levels in AD [50, 51] and potentially associated with disease progression. Thus, the ability of [^18^F]GPR39-2.1 to sensitively detect these changes highlights its potential as a noninvasive biomarker for monitoring GPR39 dysregulation and its contribution to AD pathophysiology. Importantly, these imaging data align with the emerging biological significance of GPR39 as a regulator of both vascular and neuronal homeostasis as it can quantify the GPR39 levels in these pathological conditions. Biodistribution studies revealed favorable pharmacokinetics characterized by renal clearance, low nonspecific binding, and minimal *in vivo* defluorination, which are essential attributes for high-quality PET imaging. Reduced brain uptake of [^18^F]GPR39-2.1 in biodistribution studies (90% in KO mice, \*\*\**p*<0.0002 and 77% in 5xFAD transgenic mice, \*\**p*<0.002) were significant and complementing PET imaging findings. Autoradiography provides orthogonal confirmation at the tissue level, showing strong baseline binding and substantial displacement (62-80%) by selective blocking agents. The cerebellum could be a reference region for calculating SUVR values of the PET imaging based on the autoradiography as the binding remains unchanged. Collectively, the consistent agreement across computational predictions, biochemical assays, cellular studies, and animal studies provides compelling evidence that [^18^F]GPR39-2.1 is a highly specific, stable, and translationally relevant radiotracer.

Given the growing evidence linking GPR39 to vascular inflammation, neurodegeneration, and substance-related neuroadaptations, this tracer offers a powerful, noninvasive tool for interrogating GPR39 function *in vivo* and holds considerable promise for advancing both mechanistic studies and therapeutic monitoring across neurological, cardiovascular, and neuropsychiatric diseases. Limitation of the current study includes the use of an exclusively male cohort, which does not account for potential sex-dependent differences in GPR39 expression, particularly relevant given known sexual dimorphism in AD progression and neuroinflammation. Additionally, the cross-sectional design limits insight into temporal changes in GPR39, as longitudinal dynamics relative to amyloid-β deposition and cognitive decline were not assessed. Furthermore, species-specific differences in receptor distribution and metabolism limit direct clinical translation from rodent models. Future studies will address these gaps by incorporating sex-balanced cohorts, conducting longitudinal PET imaging to map disease progression, and evaluating tracer performance in non-human primates to better approximate human neurobiology. In parallel, expanding the application of [^18^F]GPR39-2.1 to other GPR39-associated conditions, including alcohol use disorder and cardiovascular diseases, will further establish its utility beyond neurodegeneration.

This study presents the successful development and validation of [^18^F]GPR39-2.1 as a first [^18^F] radiotracer for imaging GPR39 *in vivo*. Through *in silico* studies, radiochemistry, and *in vitro* and *in vivo* biological validation, it was demonstrated that [^18^F]GPR39-2.1 possesses characteristics required for CNS molecular imaging of GPR39, including efficient BBB penetration, high binding affinity, and favorable pharmacokinetics. Importantly, the radiotracer exhibits strong receptor specificity, as evidenced by consistent reductions in uptake across pharmacological blockade, genetic knockout, and disease models. The ability of [^18^F]GPR39-2.1 to detect alterations in GPR39 levels in the 5xFAD model highlights its potential utility as a marker for neurodegenerative disease processes. Despite these promising findings, further studies are required to evaluate tracer performance in higher species and to assess its translational potential in clinical settings. Expanding its application to other GPR39-related pathologies, including cardiovascular and neuropsychiatric disorders, will also be essential to fully realize its diagnostic and therapeutic monitoring capabilities. Collectively, [^18^F]GPR39-2.1 represents a powerful and versatile tool for noninvasive imaging of GPR39, offering new opportunities to advance our understanding of its role in health and disease and to facilitate the development of targeted interventions.

## 4. Methods

### 4.1 In silico analysis

For molecular docking studies, human GPR39 protein structure (Uniprot No. O43194) was obtained from AlphaFold Protein Structure database [42]. A structure data file of lead compound (GTPL7803; Compound CID: 86277841) was retrieved from PubChem database. The lead molecule was subsequently derived to generate GPR39-2.1, GPR39-2.2, and GPR39-2.3 using BIOVIA Discovery Studio Visualizer. Molecular docking studies of GPR39 with lead molecule and its derivatives (GPR39-2.1, GPR39-2.2, and GPR39-2.3) were carried out in extra precision mode using the Glide module of the Schrodinger software suite (version 2024-2) [52]. The pharmacokinetic properties of the lead and designed GPR39 ligands were evaluated using the BOILED-Egg model and SwissADME to predict their ability to permeate the BBB [53, 54].

### 4.2 Radiochemistry

#### 4.2.1 Synthesis of [^18^F]GPR39 tracers

All radiochemical synthesis were automated on the GE TRACERLab FX_FN_ module (**Scheme 2A**). The [^18^F]F^−^ ion was produced via the ^18^O(*p,n*)^18^F nuclear reaction by irradiating [^18^O]H_2_O with protons accelerated by a cyclotron (GE PETtrace). [^18^F]F^−^ was transferred over a QMA SPE cartridge and eluted into the reaction vial with an acetonitrile solution (ACN, 1 mL) containing Kryptofix (K_2.2.2._, 50 mg), potassium carbonate (K_2_CO_3_, 5 mg) and water (100 µL). An activated [^18^F]KF complex was produced through azeotropic drying, with an additional acetonitrile (0.5 mL) addition at 140 °C under vacuum and Ar gas flow. A solution of precursor (**5** (2 mg) in DMF (1 mL)); or **10** (1 mg) in DMSO (1 mL); or **15** (1mg) in DMSO (1 mL)) was added to the activated [^18^F]KF complex. The reaction mixture was heated to 140 °C for 25 min ([^18^F]**6**); or 120 °C for 10 min ([^18^F]**11** or [^18^F]**16**). After cooling the reaction mixture to 40 °C, a quenching solution of 1:1 ACN:H_2_O (3 mL) and loaded into the semipreparative HPLC loop. A Luna^®^ 10 μm C18 100 Å LC Column 250 × 10 mm column with an isocratic flow rate of 4 mL/min and a UV = 254 nm was used for all three compounds. A mobile phase of ACN:H_2_O (70/30) was used for compound [^18^F]**6** (**t_R_** = 24.5–27.5 min). Where a mobile phase of ACN:H_2_O (80/20) was used for both [^18^F]**11** and [^18^F]**16**, both eluting at *t_R_* = 13.0-13.1 min. The collected fraction for each radiotracer was diluted in the collection flask containing water (25 mL) prior to loading onto an activated C18 Sep-Pak, followed by a water (15 mL) wash and ethanol (1.5 mL) elution. Saline (10 mL) was added to the eluted fraction to obtain an isotonic solution ready for injection.

#### 4.2.2 Quality control of [^18^F]GPR39 tracers

Chemical and radiochemical purity were assessed using an analytical HPLC (Prodigy™ 5 μm ODS-3 100 Å, LC Column 250 × 4.6 mm) at a flow rate of 1.0 mL/min, with a mobile phase of ACN:H_2_O (95/5, v/v) for [^18^F]**6** (*t_R_* = 6.9–7.4 min) and ACN:H_2_O (90/10, v/v) for both [^18^F]**11** (*t_R_*= 4.9–5.2 min) and [^18^F]**16** (*t_R_* = 6.2–6.4 min). A UV wavelength of 254 nm was used for all compounds. Respective non-radioactive compounds were used in co-injections for reference.

### 4.3 Serum stability

*In vitro* serum stability of [^18^F]GPR39-2.1, [^18^F]GPR39-2.2, and [^18^F]GPR39-2.3 were evaluated in human serum. Each radiotracer (37-40 MBq) was added to human serum (1 mL total volume) and incubated at 37 °C. Aliquots at different time points were collected at 0, 30, 60, 90 and 120 min. Samples were analyzed by radio-HPLC to determine radiochemical purity and assess the formation of radiolabeled metabolites [55]. All three tracers were run on the same Phenomenex HPLC column, 250 x 4.6 mm; 5 μm and UV_max_ = 254 nm; flow rate = 1.0 mL/min. Mobile phase = 95% acetonitrile and 5% water for [^18^F]GPR39-2.1 and mobile phase = 90% acetonitrile and 10% water for [^18^F]GPR39-2.2 and [^18^F]GPR39-2.3.

### 4.4 Inositol monophosphate assay

GPR39 receptor activity was measured using an inositol monophosphate (IP1) accumulation assay. MC38 cells expressing human GPR39 were cultured under standard conditions (37 °C, 5% CO₂) and 10,000 cells were seeded into 96-well plates 24 h prior to the experiment. On the day of the assay, the culture medium was replaced with assay buffer containing lithium chloride (LiCl) to inhibit inositol phosphate degradation and allow IP1 accumulation. The GPR39-2.1 compound was prepared in serial dilutions spanning a concentration range of 0.5 nM to 1 mM and added to the cells in triplicate wells. Following compound treatment, cells were incubated at 37 °C for 2 h to facilitate receptor activation and downstream signaling. IP1 levels were subsequently measured using a HTRF IP-One Gq detection kit (Revvity, Waltham, MA). Fluorescence signals were detected using a compatible plate reader and expressed as the ratio of emission intensities at 665 nm and 620 nm. Dose–response curves were generated by plotting IP1 accumulation against the logarithm of compound concentration. Nonlinear regression analysis was performed using GraphPad Prism to determine pharmacological parameters, including EC_50_ and maximal response (E_max_).

### 4.5 Cell uptake study

Human breast cancer (BT-20), murine colon cancer (MC38), human prostate cancer (PC3), and human glioblastoma (U87-MG) cell lines were used to evaluate cellular uptake of [^18^F]GPR39-2.1. All cell lines were cultured under standard conditions in appropriate growth media supplemented with fetal bovine serum and maintained at 37 °C in a humidified atmosphere containing 5% CO_2_. Cells were harvested at approximately 70–80% confluency for uptake experiments. Cells (∼ 1 million/well) were seeded in 6-well plates and allowed to adhere overnight. Time-dependent cellular uptake of [^18^F]GPR39-2.1 was assessed in PC3 and U87-MG cell lines. On the day of the experiment, cells were incubated with media containing [^18^F]GPR39-2.1 (0.037 MBq/well) at 37 °C for different time intervals (up to 120 min). At specified time points (5, 15, 30, 60, and 120 min), the medium was removed, and cells were washed with phosphate-buffered saline (PBS) to remove unbound radiotracer. Following washing, cells were lysed by adding 0.1 N NaOH, and the retained radioactivity was quantified using a gamma counter. To evaluate the specificity of [^18^F]GPR39-2.1 binding toward GPR39, competitive inhibition studies were performed. On the day of the experiment, cells were pre-incubated with known GPR39 agonists, TC-G1008 and TM-N1324, at a concentration of 100 µM for 1 h prior to the addition of the radiotracer. After pre-treatment, [^18^F]GPR39-2.1 was added, and the uptake assay was conducted under identical conditions as described above. The uptake values were normalized to total protein content measured by BCA protein estimation kit.

### 4.6 Animal sources and ethical approvals

BALB/c (aged 6 months, male), 5xFAD mice (aged 18 months, male), Wistar IGS rats (aged 6 months, male) were procured from the Jackson Laboratory (Bar Harbor, ME, USA). GPR39 knockout (KO) (aged 18 months, male) mice were obtained from Dr. Anthony Barnes, from Oregan Health & Science University, Portland, OR [56]. All procedures involving mice were reviewed and approved by the Wake Forest University Institutional Animal Care and Use Committee (protocols: A23-144 and A25-026), with measures taken to minimize any distress or suffering experienced by the animals.

### 4.7 In vivo microPET/CT imaging

To evaluate the *in vivo* imaging potential of [^18^F]GPR39-2.1, PET/CT studies were conducted using a small-animal microPET/CT system (TriFoil Imaging, Chatsworth, CA, USA). For all animal studies, prior to radiotracer administration, they were anesthetized in an induction chamber using 1.5-2.0% isoflurane in oxygen and maintained under anesthesia throughout the imaging procedure [52, 57]. Six-month-old healthy male BALB/c mice were used for baseline imaging studies. Each animal received an intravenous bolus injection of [^18^F]GPR39-2.1 (∼5.5 ± 0.5 MBq) via the tail vein. For TAC studies, PET imaging was performed over 0-120 min. For all other imaging studies, dynamic PET data were acquired in list mode for 60 min, with the whole-body positioned within the field of view of the scanner. CT scans were subsequently obtained for anatomical reference, and PET images were automatically co-registered with the corresponding CT images. To assess the specificity of [^18^F]GPR39-2.1 binding to GPR39 *in vivo*, blocking studies were performed using the known GPR39 agonist TC-G1008. BALB/c mice (n = 5/group) were pre-treated with TC-G1008 (20 mg/kg, IP) one hour prior to radiotracer administration. PET imaging was conducted under both baseline and blocking conditions to evaluate receptor-specific binding. To further investigate brain-specific uptake and regional distribution, PET imaging studies were also conducted in higher rodent models such as Wistar IGS rats (n = 4/group). Animals were imaged under baseline and blockade with TC-G1008 (10 mg/kg, i.p., administered 1 h prior to tracer injection). To confirm the target specificity of [^18^F]GPR39-2.1, comparative PET imaging studies were performed in wild-type (n = 5) and GPR39 KO mice (n = 4). Differences in tracer uptake between these groups were used to validate GPR39-specific binding. To evaluate the applicability of [^18^F]GPR39-2.1 in disease conditions, PET imaging studies were conducted in the AD mouse models (5xFAD). Both transgenic and age-matched wild-type mice (n = 5/group) were imaged under identical conditions, and tracer uptake was compared to assess potential alterations in GPR39 levels associated with disease pathology.

PET data were reconstructed using a three-dimensional ordered subset expectation maximization (3D-OSEM) algorithm provided by TriFoil Imaging. Whole body PET/CT images were generated using PMOD software (version 4.3; PMOD Technologies LLC, Zurich, Switzerland). After removing the animal bed from the CT dataset, the bed-stripped CT volume was used to generate 2D PET/CT fusion images in PBAS, improving visualization of tracer distribution. Quantitative image analysis was performed using PMOD software (version 4.3; PMOD Technologies LLC, Zurich, Switzerland). Regions of interest (ROIs) were automatically delineated, and tracer uptake was quantified. TACs were generated for the whole brain (0-120 min). Standardized uptake values (SUVs) were calculated and averaged over the dynamic acquisition period (60 min) according to established protocols [58–60].

### 4.8 Tissue biodistribution studies

Tissue biodistribution of [^18^F]GPR39-2.1 was evaluated in male BALB/c mice (n = 5/time point) at 5, 30, 60, 90, and 120 min post-injection, following previously established protocol [61]. Biodistribution studies were also performed in Wistar IGS rats (n = 4/group) following PET imaging studies under both baseline and blocking conditions using TC-G1008. To further validate tracer specificity, *ex vivo* biodistribution analyses were performed in the same animals used for imaging studies, including wild type (n = 5) and GPR39 KO mice (n = 4) and 5xFAD wild type and transgenic mice (n = 5/group). All collected tissues were weighed and measured for radioactivity using a gamma counter, with a diluted aliquot of the injected radiotracer used as a calibration standard [52, 61, 62]. The uptake of [^18^F]GPR39-2.1 in each tissue was expressed as the percentage of injected dose per gram of tissue (%ID/g), enabling quantitative comparison of tracer distribution across organs, time points, and experimental conditions.

### 4.9 Autoradiography studies

*In vitro* autoradiography was performed to assess the regional distribution and specific binding of [^18^F]GPR39-2.1 in mouse brain tissue sections following established protocols [58, 62]. In brief, brain tissues from BALB/c mice (n = 4) were rapidly harvested and cryosectioned into 20 µm thick sections using a cryostat. Each tissue section was incubated with [^18^F]GPR39-2.1 (37 KBq/mL in 1× PBS, pH 7.4) at room temperature for 1 h. For blocking experiments, tissue sections were pre-incubated with the GPR39 agonist TC-G1008 (100 µM) for 1 h prior to radiotracer exposure to evaluate receptor-specific binding. Following incubation, sections were washed four times with PBS (3 min per wash) to remove unbound radiotracer. The slides were then air-dried and exposed overnight to a BAS-IP-SR phosphor imaging plate at −20 °C. The exposed plates were subsequently scanned using a GE Amersham Typhoon scanner with a spatial resolution of 25 µm. Autoradiographic images were analyzed using ImageQuant TL 8.2 software, and tracer binding was quantified using MCID Core 7.1 software. Specific binding was calculated by comparing baseline and blocking conditions and expressed as photostimulated luminescence per unit area (PSL/mm²).

## Author Contributions

All authors contributed to the study conception and design. Material preparation, data collection and analysis were performed by **Bhuvanachandra Bhoopal, Ryan W Fitzgerald and Kiran K Solingapuram Sai**. The first draft of the manuscript was written by **Bhuvanachandra Bhoopal** and **Ryan W Fitzgerald** and all authors commented on previous versions of the manuscript. All authors read and approved the final manuscript.

## Data Availability Statement

All data generated or analyzed during this study are included in this published article and/or its Supplementary Information files.

