## supplemental for "First-in-class [^18^F]-PET radiotracers for GPR39: A platform for noninvasive target engagement imaging *in vivo*"

### 1. Chemical synthesis

#### 1.1. Synthesis of 6-chloro-N-(2,4-dichlorobenzyl)-N,2-dimethylpyrimidin-4-amine 3:

To a stirred solution of 4,6-dichloro-2-methylpyrimidine **1** (8 g, 49.079 mmol, 1.0 eq.) in anhydrous THF (50 mL), (2,4-dichlorophenyl)-N-methylmethanamine **2** (7.9 g, 41.717 mmol, 0.85 eq.) and DIPEA (17 mL, 2.0 eq.) were added at room temperature, and the reaction mixture was stirred at room temperature overnight. Progress of the reaction was monitored by TLC, after completion of the reaction, the reaction mixture was quenched and diluted with water (100 mL), and the product was extracted with DCM (2 × 100 mL). The combined organic layers were washed with water (100 mL), brine (2 x 50 mL) and dried over sodium sulfate and concentrated under reduced pressure. The crude compound was purified by silica gel column chromatography using with elution of MeOH and DCM solvent mixture to afford 6-chloro-N-(2,4-dichlorobenzyl)-N,2-dimethylpyrimidin-4-amine **3** (6 g, yield: 38.7%) as an off white solid.

^1^H-NMR (CDCl_3_, 400MHz): δ 2.48 (s, 3H), 3.06 (bs, 3H), 4.86 (bs, 2H), 6.26 (s, 1H), 6.99 (d, *J* = 6.4 Hz, 1H), 7.19 (dd, *J* = 6.4 Hz, 1.2 Hz, 1H), 7.42 (s, 1H).

#### 1.2. Synthesis of N-(2,4-dichlorobenzyl)-N,2-dimethyl-6-(6-nitropyridin-3-yl)pyrimidin-4-amine 5:

To a stirred solution of 6-chloro-N-(2,4-dichlorobenzyl)-N,2-dimethylpyrimidin-4-amine **3** (1.1 g, 3.473 mmol, 1.0 eq.) and 5-(4,4,5,5-tetramethyl-1,3,2-dioxaborolan-2-yl)-2-nitropyridine **4** (0.95 g, 3.821 mmol, 1.1 eq.) in anhydrous THF (20 mL), was added sodium carbonate (1.1 g, 3.0 eq.) and the mixture was degassed with argon for 10 min. Then, Pd(dppf)Cl_2_.DCM (0.52 g, 0.694 mmol, 0.2 eq.) was added and the reaction mixture was heated to 110˚C and stirred for 90 min. Progress of the reaction was monitored by TLC, after completion of the reaction, cooled to ambient temperature, filtered through celite pad and filtrate was concentrated under reduced pressure. The crude compound was purified by silica gel column chromatography using with elution of EtOAc and hexane solvent mixture to afford N-(2,4-dichlorobenzyl)-N,2-dimethyl-6-(6-nitropyridin-3-yl)pyrimidin-4-amine **5** (0.6 g, 1.48 mmol, yield: 42%) as an off white solid.

^1^H-NMR (CDCl_3_, 400MHz): δ 2.59 (s, 3H), 3.20 (bs, 3H), 4.95 (bs, 2H), 6.65 (s, 1H), 7.07 ( d, *J* = 6 Hz, 1H), 7.19 (dd, *J* = 6.8 Hz, 1.2 Hz, 1H), 7.43 (s, 1H), 8.32 ( d, *J* = 6.8 Hz, 1H), 8.62 (d, *J* = 6.0 Hz, 1H), 9.09 (s, 1H). ES-MS: m/z 404.26 [M+H]^+^. HPLC purity: 99.11 % (RT: 29.693 min).

#### 1.3. Synthesis of N-(2,4-dichlorobenzyl)-6-(6-fluoropyridin-3-yl)-N,2-dimethylpyrimidin-4-amine 6:

To a stirred solution of N-(2,4-dichlorobenzyl)-N,2-dimethyl-6-(6-nitropyridin-3-yl)pyrimidin-4-amine **5** (0.4 g, 0.989 mmol, 1.0 eq.) in dimethyl sulfoxide (8 mL), tetramethylammonium fluoride (1.84 g, 19.789 mmol, 20 eq.) was added. The reaction mixture was heated to 80˚C and continued stirring for 8 h. Progresses of the reaction was monitored by TLC, after completion of the reaction, cooled to ambient temperature, diluted with ethyl acetate (20 mL) and filtered. The filtrate was washed with water (10 mL) dried over anhydrous sodium sulfate and filtered and concentrated under reduced pressure. The crude compound was purified by silica gel column chromatography using with elution of EtOAc and hexane solvent mixture to afford N-(2,4-dichlorobenzyl)-6-(6-fluoropyridin-3-yl)-N,2-dimethylpyrimidin-4-amine **6** (0.25 g, 0.662 mmol, yield: 67%) as a white solid.

^1^H-NMR (CDCl_3_, 400MHz): δ 2.57 (s, 3H), 3.16 (bs, 3H), 4.93 (bs, 2H), 6.55 (s, 1H), 7.01 (d, *J* = 5.6 Hz, 1H), 7.06 (d, *J* = 6.4 Hz, 1H), 7.18 (d, *J* = 6.4 Hz, 1H), 7.43 (s, 1H), 8.39 (t, *J* = 6.6 Hz, 1H), 8.72 (s, 1H). ES-MS: m/z 377.40 [M+H]^+^. HPLC purity: 99.87 % (RT: 9.45 min).

#### 1.4. Synthesis of 2-chloro-N-(2,4-dichlorobenzyl)-N-methyl-6-(pyridin-2-yl)pyrimidin-4-amine 8:

To a stirred solution of 2,4-dichloro-6-(2-pyridinyl)pyrimidine **7** (5.0 g, 22.11 mmol, 1 eq.) and diisopropyl ethyl amine (8.57g, 66.31 mmol, 3 eq.) in acetonitrile (140.0 mL) at ambient temperature, was added 1-(2,4-dichlorophenyl)-N-methylmethanamine **2** (4.2 g, 22.09 mmol, 0.99 eq.). The reaction mass was heated to 80˚C and stirred for 16 h. Progress of the reaction was monitored by TLC. The reaction mass was allowed to room temperature and concentrated under reduced pressure. The resulting crude compound was purified by silica gel column chromatography elution of ethyl acetate and hexane mixture to afford 2-chloro-N-(2,4-dichlorobenzyl)-N-methyl-6-(pyridin-2-yl)pyrimidin-4-amine **8** (5.0 g, 13.16 mmol, yield: 59 %).

^1^H-NMR (CDCl_3_, 400MHz): δ 3.19 (s, 3H), 4.97 (s, 2H), 7.09 (s, 1H), 7.20 ( dd, *J* = 8.0 Hz, 1.6 Hz, 1H), 7.39 (t, *J* = 6.8 Hz, 1H), 7.43 (d, *J* = 1.6 Hz, 1H), 7.55 (bs, 1H), 7.85 (t, *J* = 8.0 Hz, 1H), 8.41 (d, *J* = 8.0 Hz, 1H), 8.65 (bs, 1H).

#### 1.5. Synthesis of 4-((2,4-dichlorobenzyl)(methyl)amino)-N,N,N-trimethyl-6-(pyridin-2-yl)pyrimidin-2-aminium chloride 10:

A reaction mixture of 2-chloro-N-(2,4-dichlorobenzyl)-N-methyl-6-(pyridin-2-yl)pyrimidin-4-amine **8** (4.0 g, 10.53 mmol), trimethylamine **9** (13% in THF 60 mL) in seal tube was stirred at room temperature for 72 h. The reaction mixture was cooled to 0˚C and stirred for 1 h. Progress of the reaction was monitored by TLC. The resulting solids were filtered and dried to get crude product. The crude compound was taken in THF (20 mL) and the resulting suspension was stirred at room temperature for 5 min, solids were filtered and dried under vacuum to get the impure product 4-((2,4-dichlorobenzyl)(methyl)amino)-N,N,N-trimethyl-6-(pyridin-2-yl)pyrimidin-2-aminium chloride **10** (2.6 g, 5.92 mmol, yield: 56%) as an off white color solid.

^1^H-NMR (DMSO-d_6_, 400MHz): δ 3.28 (s, 3H), 3.50 (s, 9H), 5.04 (s, 2H), 7.28 (d, *J* = 8.4 Hz, 1H), 7.39 (d, *J* = 8.0 Hz, 1H), 7.62 (m, 1H), 7.70 (s, 1H), 7.77 (s, 1H), 8.06 (t, *J* = 7.6 Hz, 1H ), 8.43 (d, *J* = 7.6 Hz, 1H), 8.80 (d, *J* = 3.6 Hz, 1H). ES-MS: m/z 402.36 [M-Cl] ^+^. HPLC purity: 99.85 % (RT: 13.39 min).

#### 1.6. Synthesis of N-(2,4-dichlorobenzyl)-2-fluoro-6-(pyridin-2-yl)pyrimidin-4-amine 11:

A solution of 4,4-((2,4-dichlorobenzyl)(methyl)amino)-N,N,N-trimethyl-6-(pyridin-2-yl)pyrimidin-2-aminium chloride **10** (1.0 g, 2.47 mmol, 1 eq.), cesium fluoride (1.0 g, 6.57 mmol, 2.9 eq.), DMSO (20 ml) was stirred at 75˚C for 8 h. Progress of the reaction was monitored by TLC. The reaction mixture was cooled to room temperature, poured into ice cold water and extracted with ethyl acetate (2 x 100 mL). The combined organic layer was washed with water and dried over sodium sulfate, filtered and concentrated under reduced pressure to get crude product. The crude compound was purified by silica gel column chromatography using with elution of ethyl acetate and hexane mixture to afford N-(2,4-dichlorobenzyl)-2-fluoro-6-(pyridin-2-yl)pyrimidin-4-amine **11** (0.5 g, 1.43 mmol, yield: 58%) as an off white solid.

^1^H-NMR (CDCl_3_, 400MHz): δ 3.21 (s, 3H), 4.98 (s, 2H), 7.07 (d, *J* = 7.2 Hz, 2H), 7.19 (dd, *J* = 8.8 Hz, 2.4 Hz, 1H), 7.39 (t, *J* = 5.6 Hz, 1H), 7.43 (s, 1H), 7.54 (s, 1H), 7.84 (t, *J* = 6.8 Hz, 1H ), 8.37 (d, *J* = 8.0 Hz, 1H), 8.67 (bs, 1H). ES-MS: m/z 363.21 [M+H] ^+^. HPLC purity: 99.66 % (RT: 20.70 min).

#### 1.7. Synthesis of 2-chloro-N-(4-chlorobenzyl)-N-methyl-6-(pyridin-2-yl)pyrimidin-4-amine 14:

To a stirred solution of 2,4-dichloro-6-(2-pyridinyl)pyrimidine **12** (5.0 g, 22.11 mmol, 1 eq.), and di-isopropyl ethyl amine (8.57g, 66.31 mmol, 3 eq.) in acetonitrile (140 mL) at ambient temperature, (4-chlorophenyl)-N-methylmethanamine **13** (3.09 g, 19.85 mmol, 0.9 eq.) was added. The reaction mixture was heated 80˚C and stirred for 16 h progress of the reaction was monitored by TLC. After completion, the reaction mass was allowed to room temperature. The reaction mass was concentrated under reduced pressure. The resulting crude compound was purified by silica gel column chromatography using with elution of ethyl acetate and hexane mixture to afford 2-chloro-N-(4-chlorobenzyl)-N-methyl-6-(pyridin-2-yl)pyrimidin-4-amine **14** (5.0 g, 14.49 mmol, yield: 65%) as an off-white solid.

^1^H-NMR (CDCl_3_, 400MHz): δ 3.13 (s, 3H), 4.88 (s, 2H), 7.19 (d, *J* = 8.4 Hz, 2H), 7.31 (d, *J* = 8.0 Hz, 2H), 7.37 (dd, *J* = 6.8 Hz, 4.8 Hz, 1H), 7.53 (s, 1H), 7.83 (td, *J* = 8.0 Hz, 1.6 Hz, 1H), 8.39 (d, *J* = 8.0 Hz, 1H), 8.65 (d, *J* = 4.0 Hz, 1H).

#### 1.8. Synthesis of 4-((4-chlorobenzyl)(methyl)amino)-N,N,N-trimethyl-6-(pyridin-2-yl)pyrimidin-2-aminium chloride 15:

A reaction mixture 2-chloro-N-(4-chlorobenzyl)-N-methyl-6-(pyridin-2-yl)pyrimidin-4-amine **14** (4.0 g, 11.58 mmol), trimethylamine **9** (13% in THF 60 mL) in a sealed tube was stirred at room temperature for 72 h. Progress of the reaction was monitored by TLC. The reaction mixture was cooled to 0˚C and stirred for 1 h. The resulting solids were filtered and dried to get crude product. The crude compound was taken in THF (20 mL) and the resulting suspension was stirred at room temperature for 5 min, solids were filtered and dried under vacuum to get the pure product 4-((4-chlorobenzyl)(methyl)amino)-N,N,N-trimethyl-6-(pyridin-2-yl)pyrimidin-2-aminium chloride **15** (2.0 g, 4.95 mmol, yield: 42%) as an off white color solid.

^1^H-NMR (DMSO-d_6_, 400MHz): δ 3.30 (s, 3H), 3.88 (s, 9H), 4.92 (s, 2H), 7.13 (d, *J* = 7.6 Hz, 2H), 7.33 (d, *J* = 7.2 Hz, 2H), 7.47 (bs, 1H), 7.80 (s, 1H), 7.92 (t, *J* = 6.8 Hz, 1H ), 8.33 (d, *J* = 7.6 Hz, 1H), 8.71 (d, *J* = 3.8 Hz, 1H). ES-MS: m/z 368.35 [M-Cl] ^+^. HPLC purity: 99.75 % (RT: 12.76 min).

#### 1.9. Synthesis of N-(4-chlorobenzyl)-2-fluoro-6-(pyridin-2-yl)pyrimidin-4-amine 16:

A solution of 4-((4-chlorobenzyl)(methyl)amino)-N,N,N-trimethyl-6-(pyridin-2-yl)pyrimidin-2-aminium chloride **15** (1.0 g, 2.47 mmol), cesium fluoride (1.12 g, 7.36 mmol) in DMSO (20 mL) was stirred at 75˚C for 8 h. Progress of the reaction was monitored by TLC (10% MeOH/DCM). The reaction mixture was cooled to room temperature, poured into ice cold water and extracted with ethyl acetate (2 x 100 mL). The combined organic layer was washed with water and dried over sodium sulfate, filtered and concentrated under reduced pressure to get crude product. The crude compound was purified by silica gel column chromatography using with elution of ethyl acetate and hexane mixture to afford N-(4-chlorobenzyl)-2-fluoro-6-(pyridin-2-yl)pyrimidin-4-amine **16** (0.52 g, 1.65 mmol, yield: 66%) as an off white solid.

^1^H-NMR (CDCl_3_, 400MHz): δ 3.19 (s, 3H), 4.97 (s, 2H), 7.09 (bs, 1H), 7.20 (dd, *J =* 8.0, 1.6 Hz, 1H), 7.26 (s, 1H), 7.39 (t, J = 2.8 Hz, 1H), 7.43 (d, *J =* 1.6 Hz, 1H ), 7.56 (bs, 1H), 7.85 (t, *J* = 8.0 Hz, 1H), 8.41 (d, *J* = 8.0 Hz, 1H), 8.66 (bs, 1H). ES-MS: m/z 329.28 [M+H] ^+^. HPLC purity: 99.51 % (RT: 18.71 min).

### 1. Tables:

**Table S1.** Physiochemical properties of lead molecule and its derivatives (GPR39-2.1, GPR39-2.2, and GPR39-2.3) predicted by SwissADME.

| **Compound** | **Lipophilicity (LogP)** | **BBB permeation** | **GI absorption** |
| --- | --- | --- | --- |
| Lead molecule (GTPL7803) | 3.68 | Yes | High |
| GPR39-2.1 | 4.45 | Yes | High |
| GPR39-2.2 | 4.16 | Yes | High |
| GPR39-2.3 | 3.65 | Yes | High |

### 2. Figures:

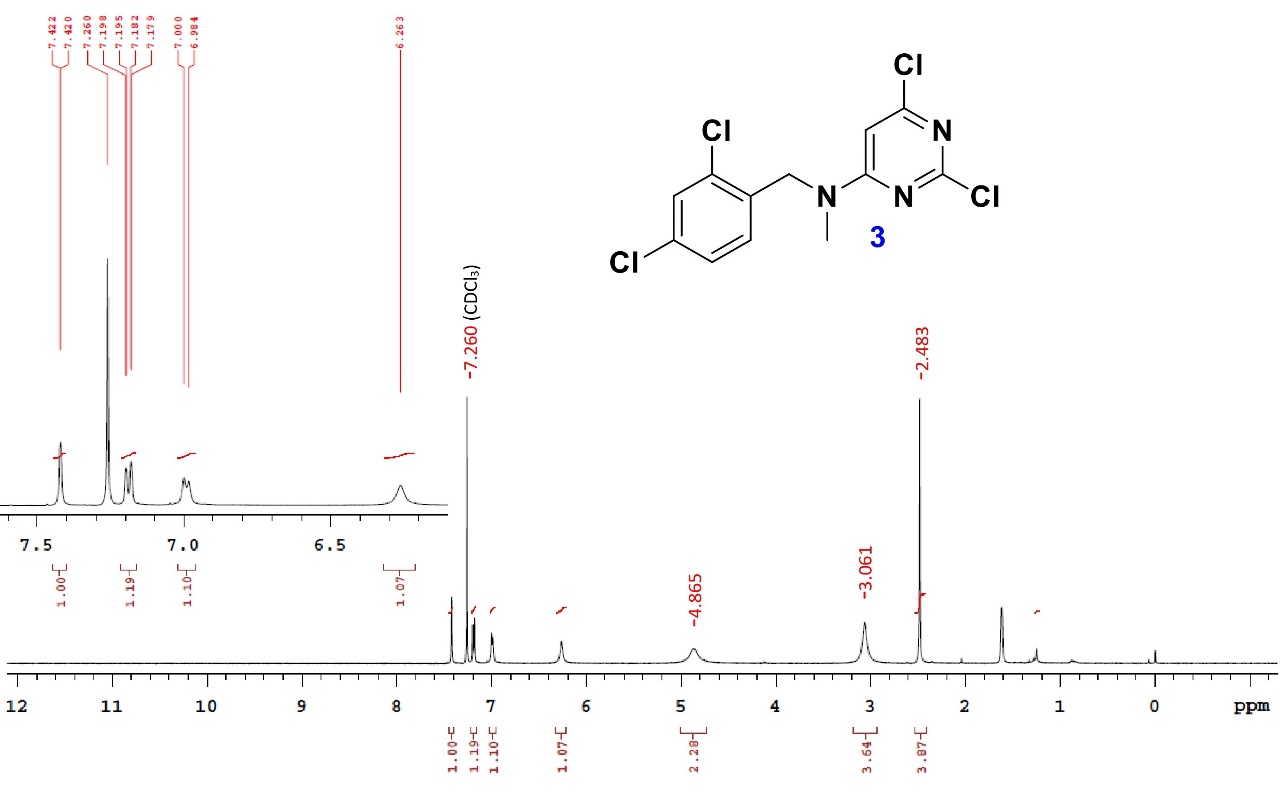

**Figure S1.** ^1^H NMR of compound **3**.

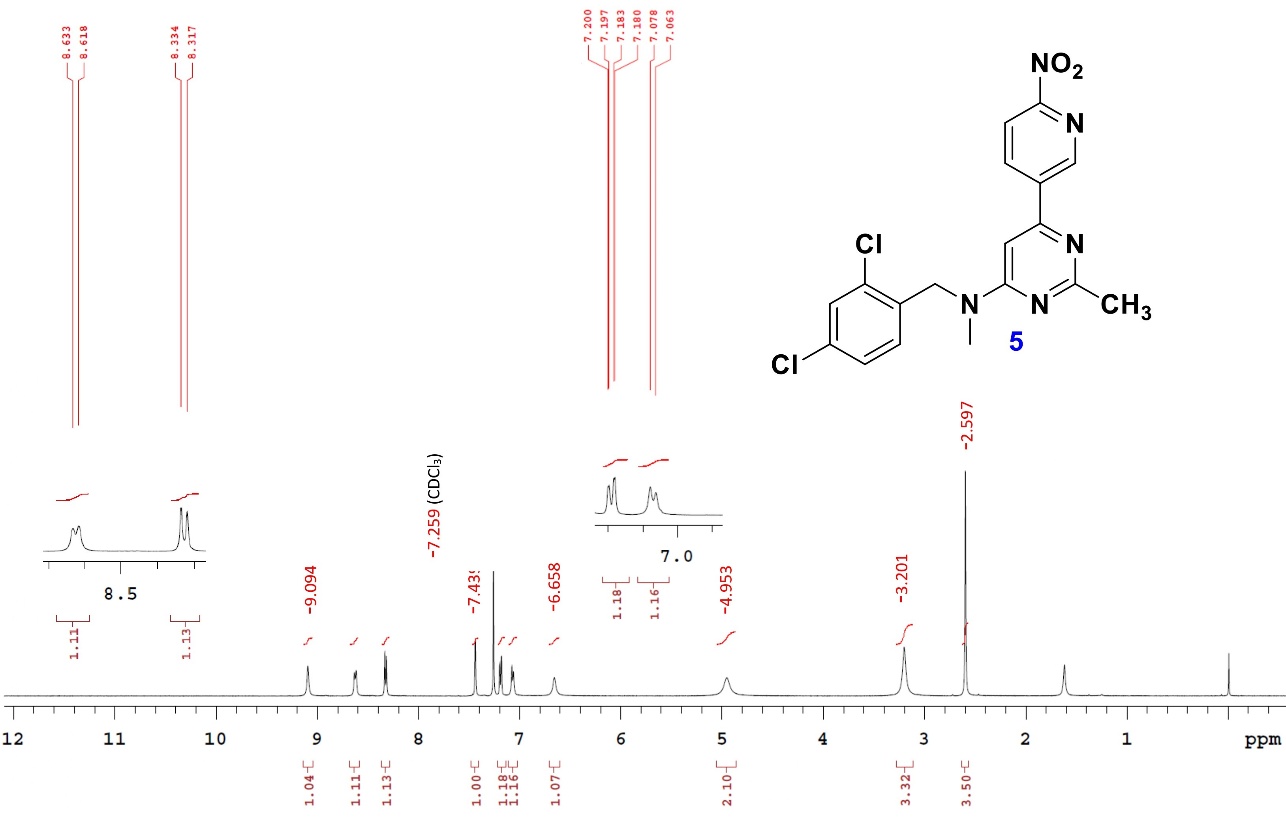

**Figure S2.** ^1^H NMR of compound **5**.

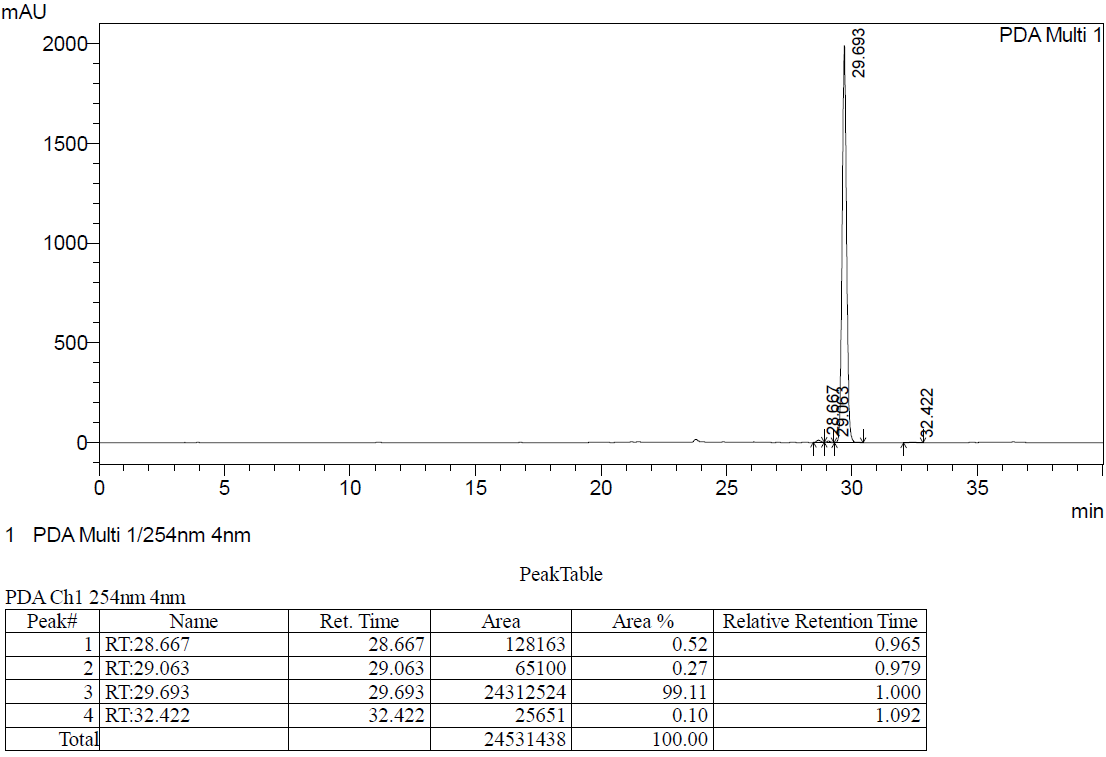

**Figure S3.** HPLC data of compound **5** (HPLC conditions: column: Inertsil ODS-3V: 250 x 4.6 mm, 5 µm; mobile phase: 50% acetonitrile in water; flow rate: 1 mL/min).

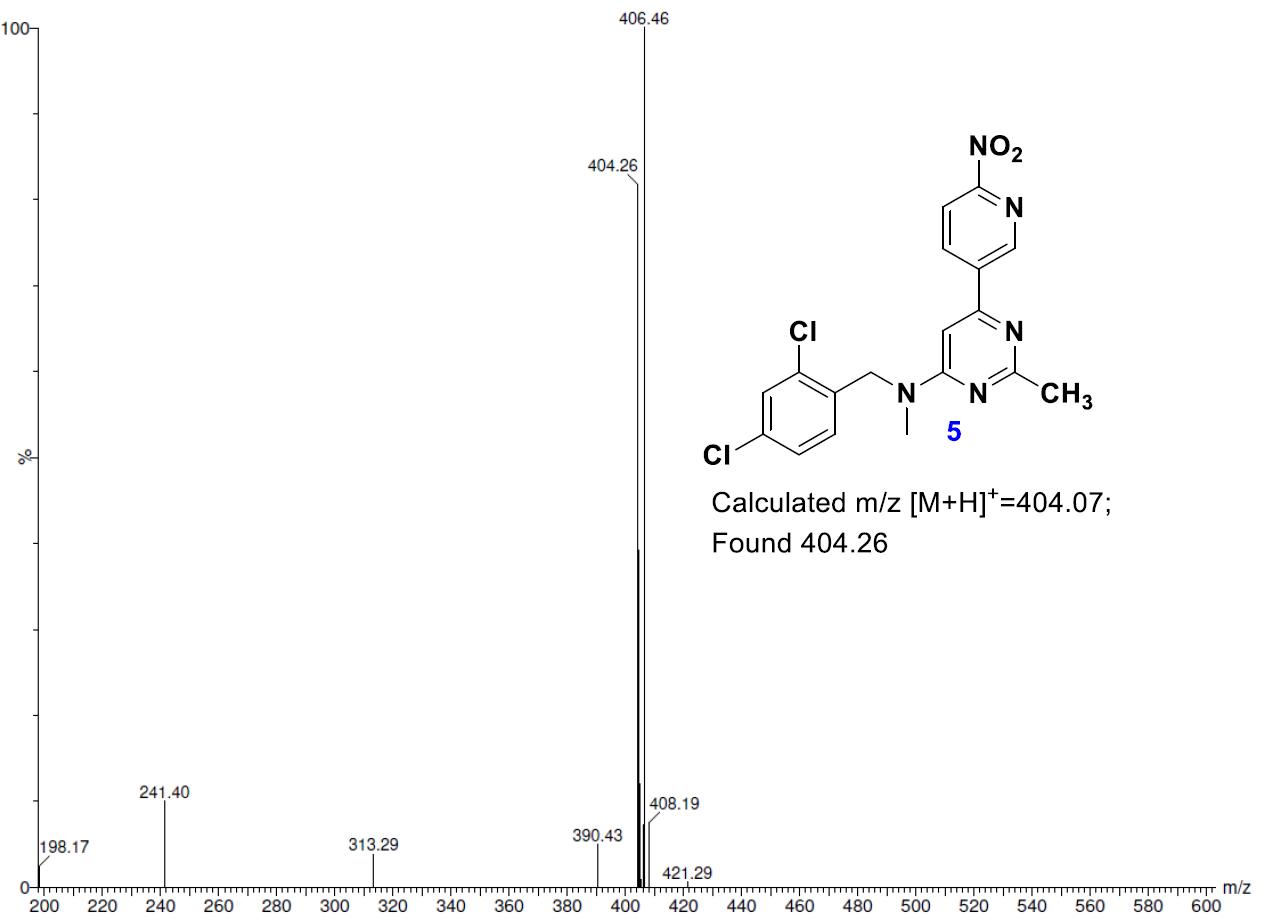

**Figure S4.** Mass spectrometry analysis of compound **5**.

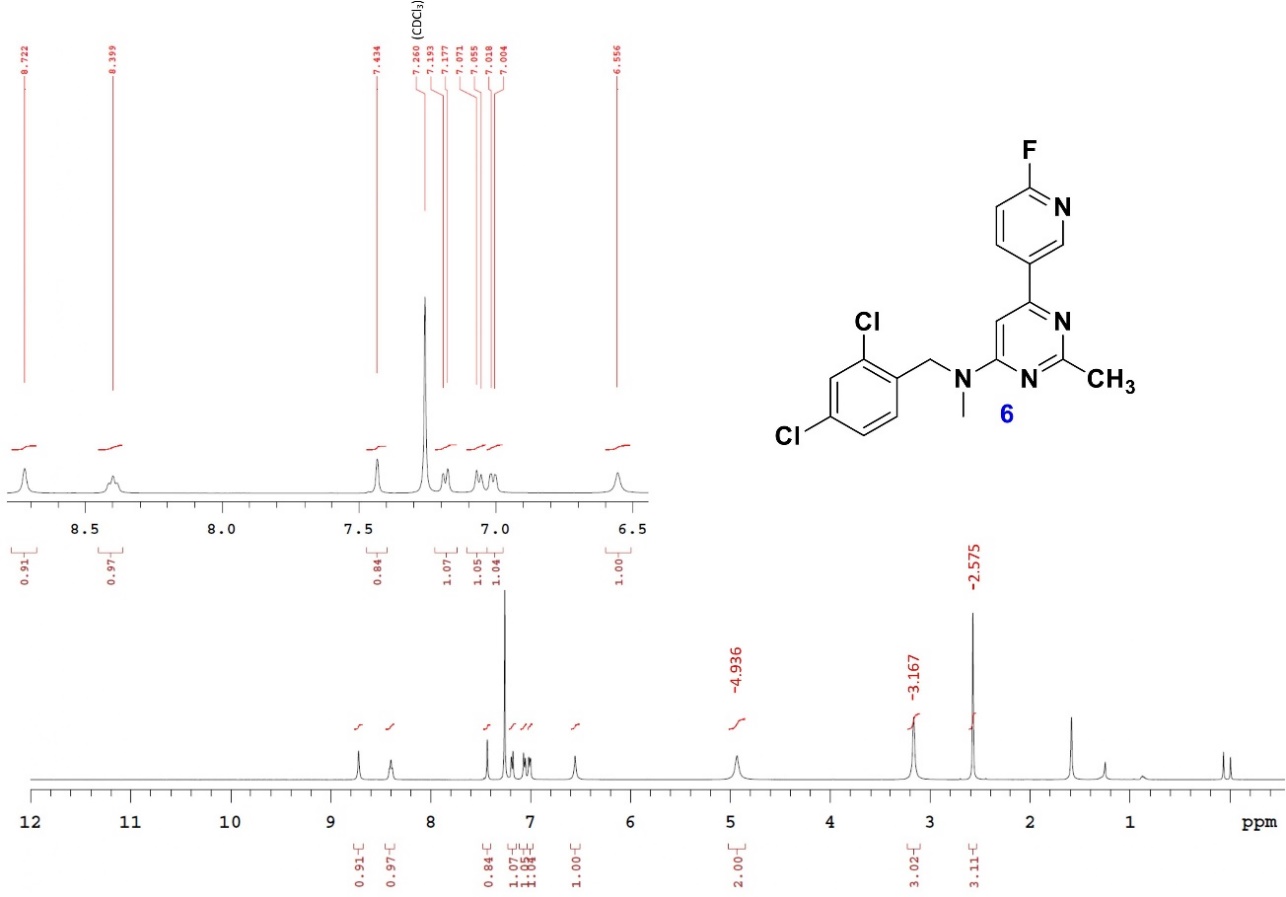

**Figure S5.** ^1^H NMR of compound **6**.

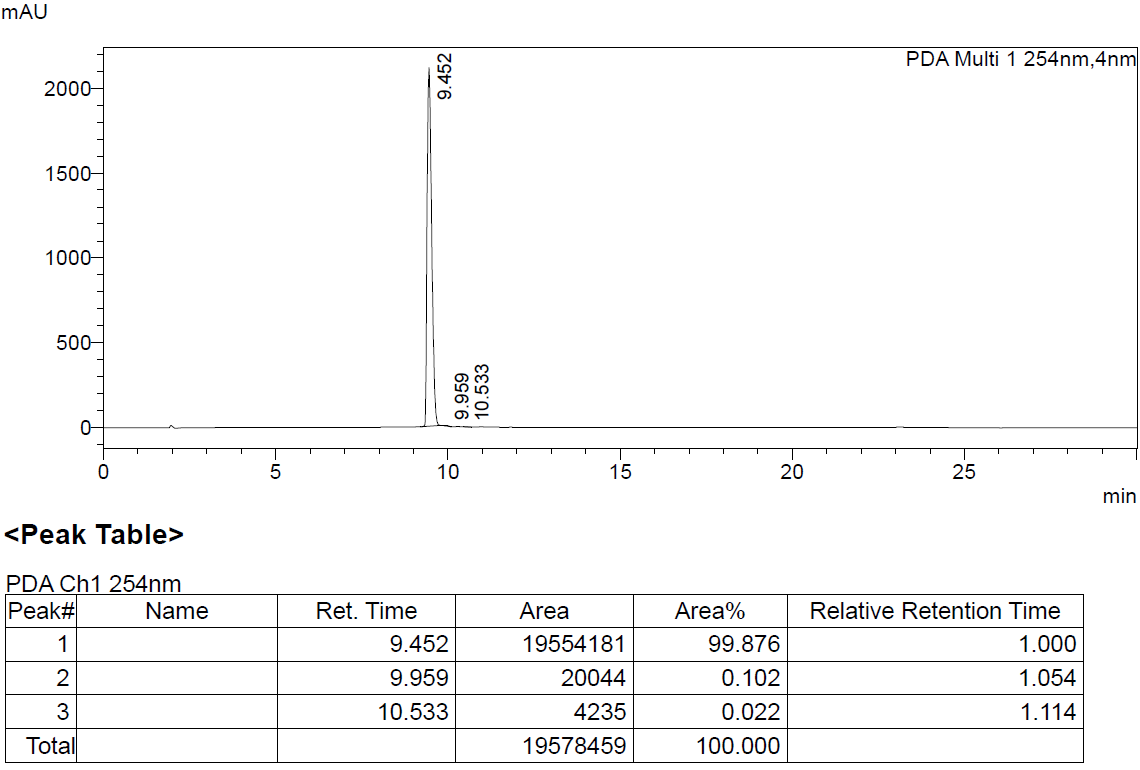

**Figure S6.** HPLC data of compound **6** (HPLC conditions: column: Kromosil C18: 150 x 4.6 mm, 5 µm; mobile phase: 50% acetonitrile in water; flow rate: 1 mL/min).

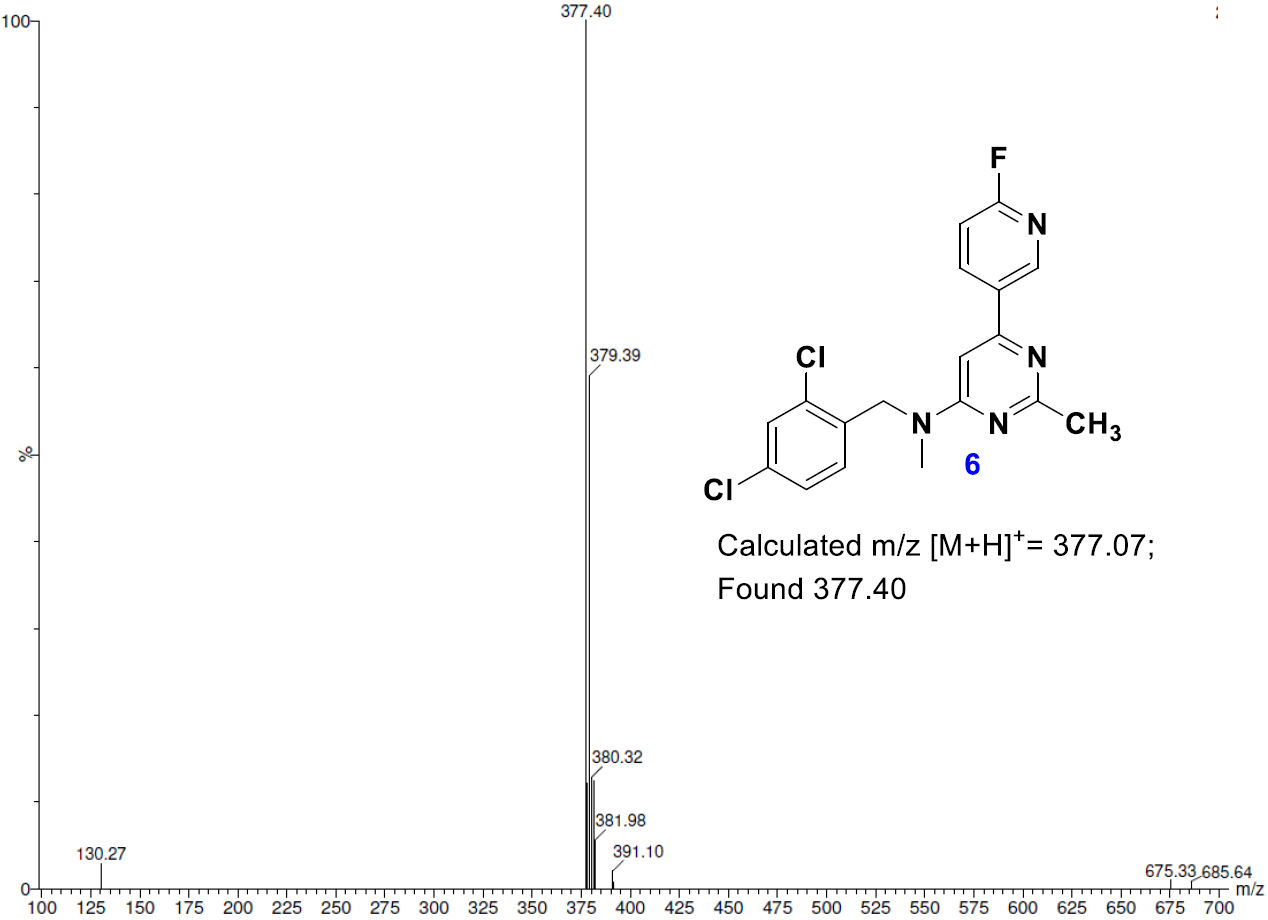

**Figure S7.** Mass spectrometry analysis of compound **6.**

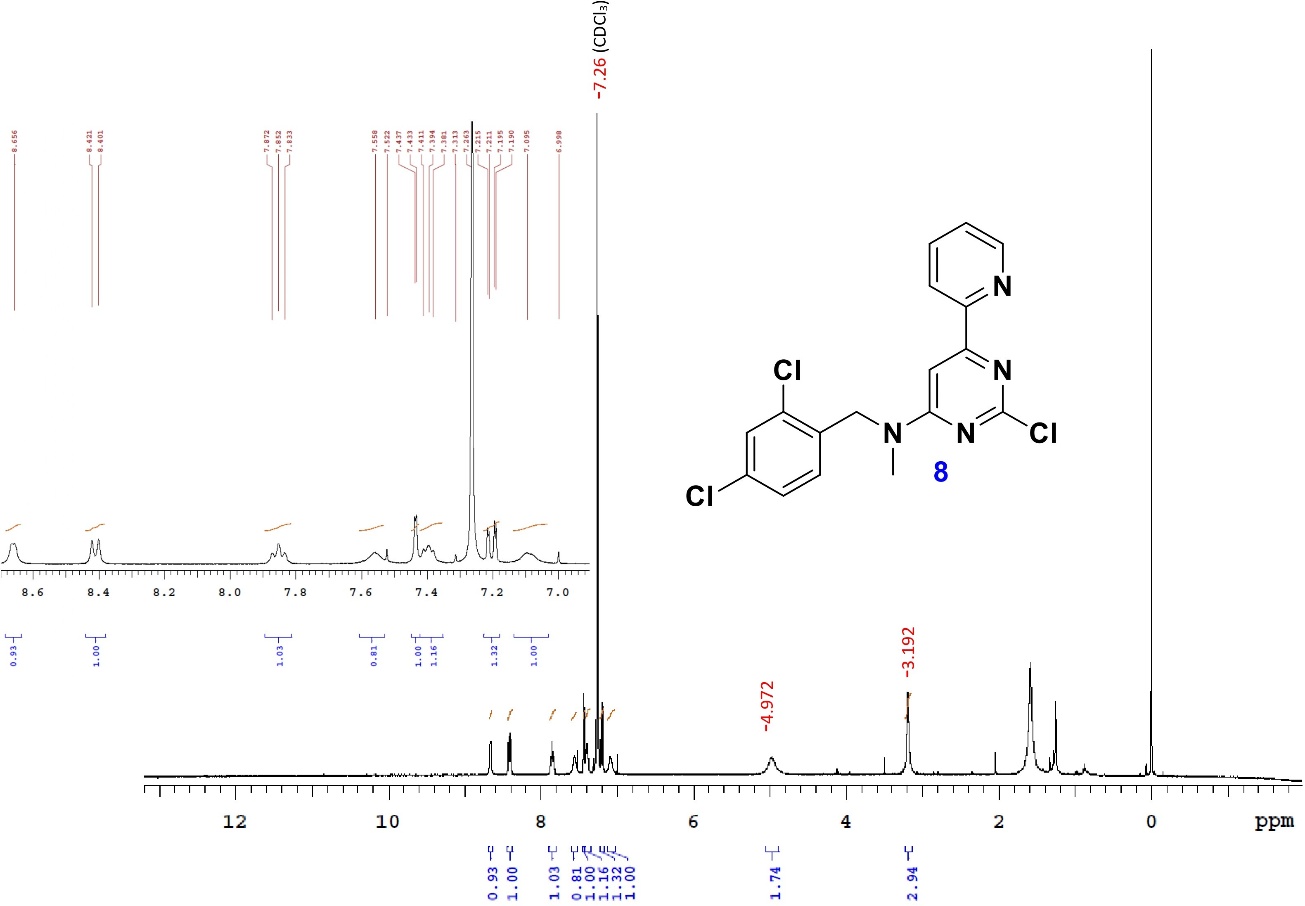

**Figure S8.** ^1^H NMR of compound **8**.

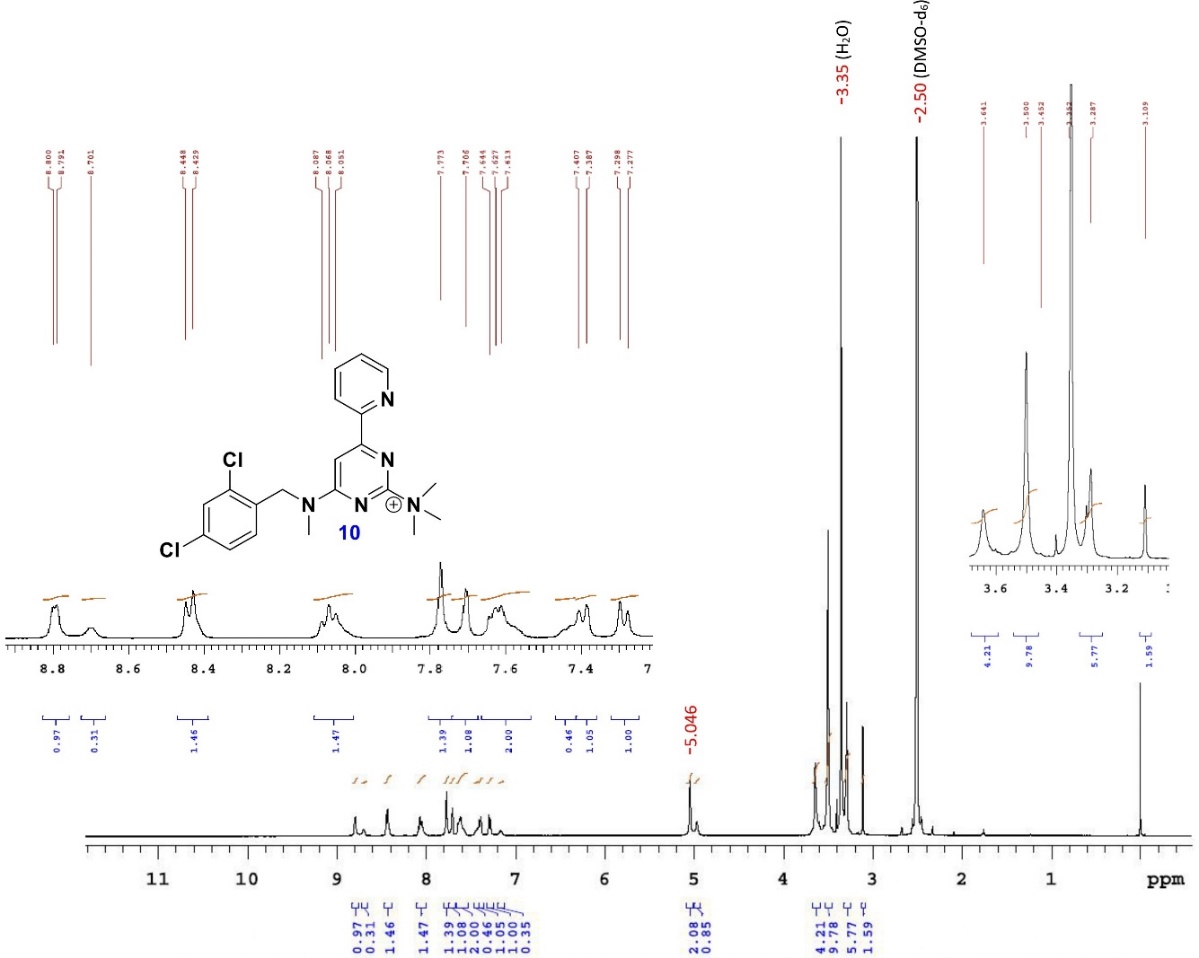

**Figure S9.** ^1^H NMR of compound **10**.

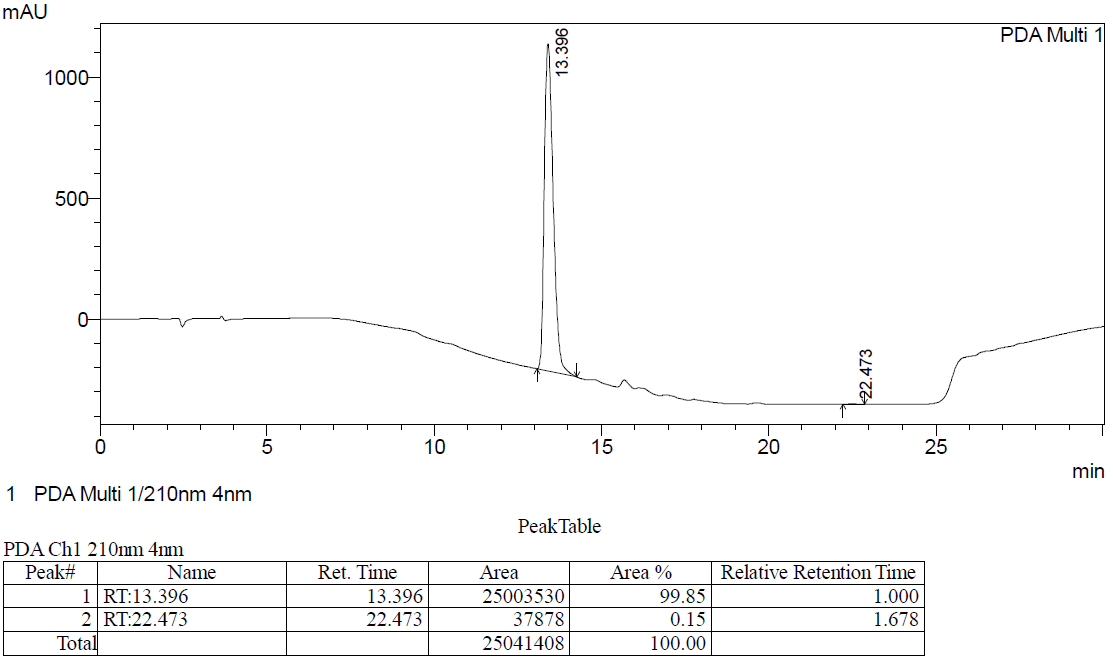

**Figure S10.** HPLC data of compound **10** (HPLC conditions: column: Inertsil ODS-3V: 250 x 4.6 mm, 5 µm; mobile phase: 50% acetonitrile in water; flow rate: 1 mL/min).

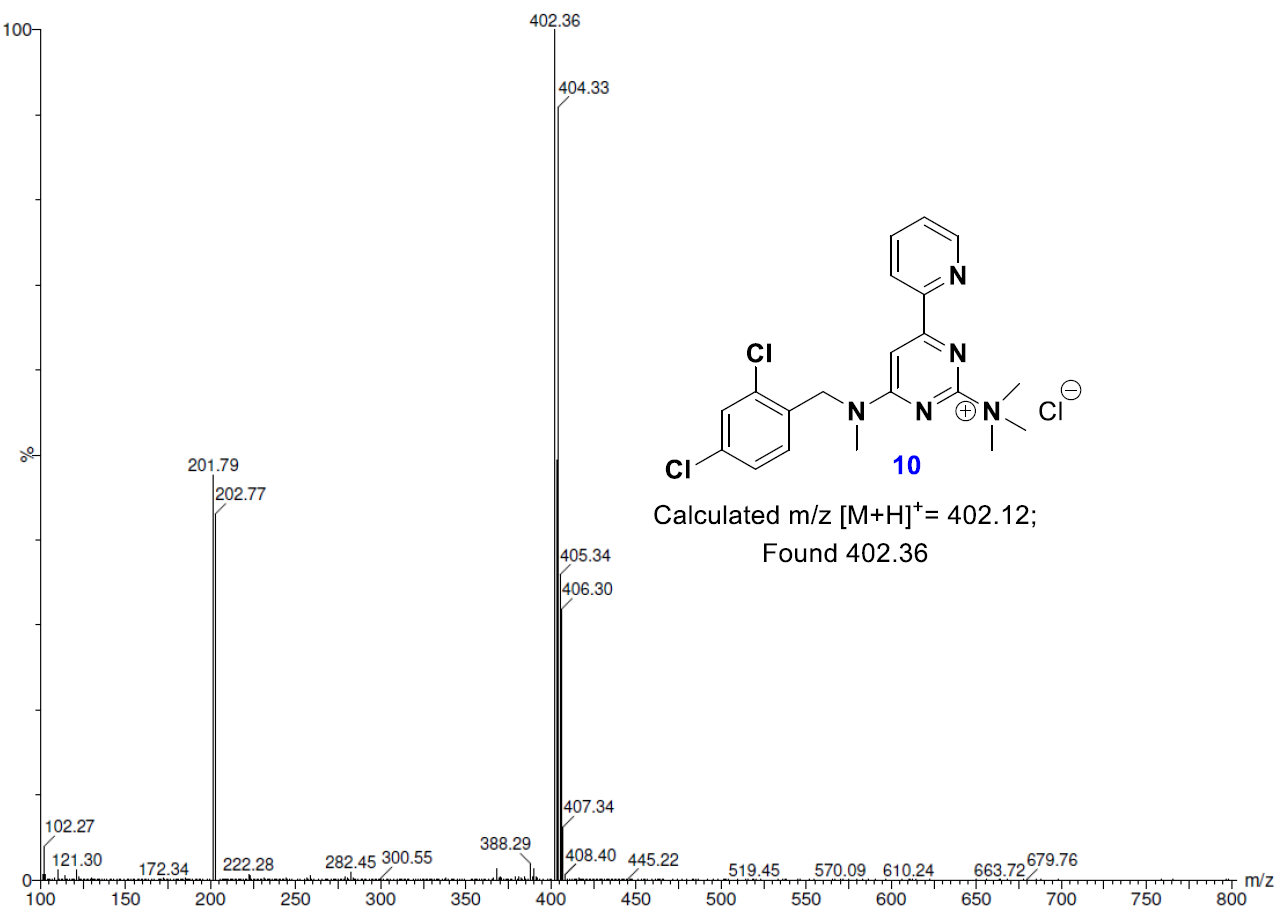

**Figure S11.** Mass spectrometry analysis of compound **10**.

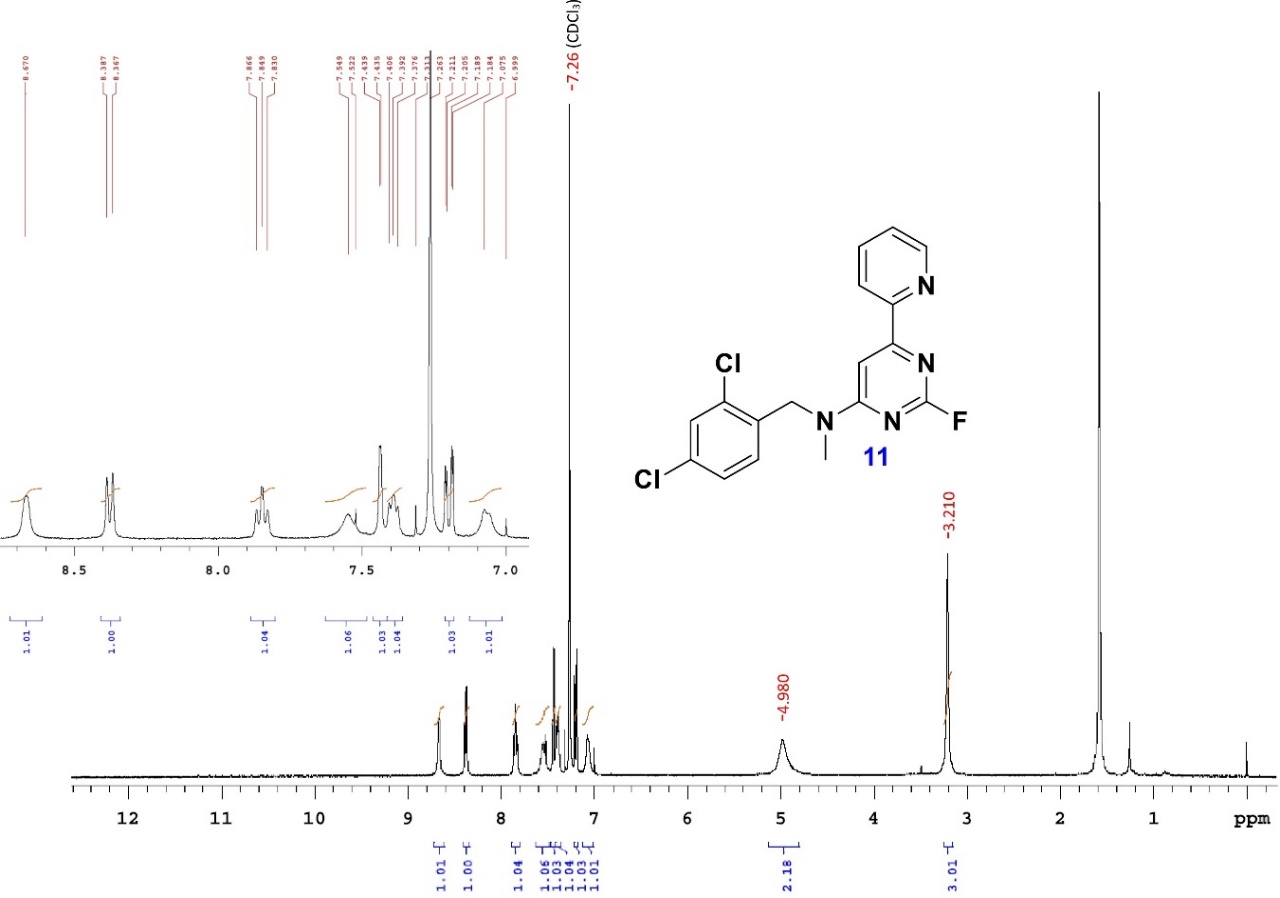

**Figure S12.** ^1^H NMR of compound **11**.

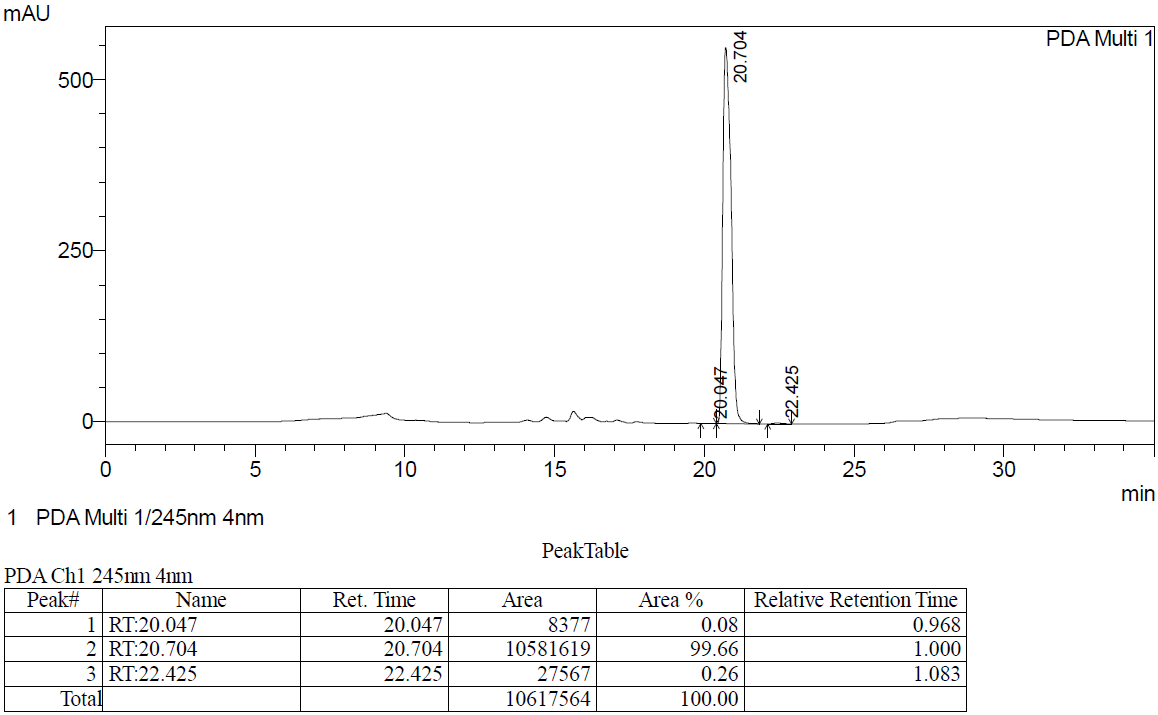

**Figure S13.** HPLC data of compound **11** (HPLC conditions: column: Inertsil ODS-3V: 250 x 4.6 mm, 5 µm; mobile phase: 50% acetonitrile in water; flow rate: 1 mL/min).

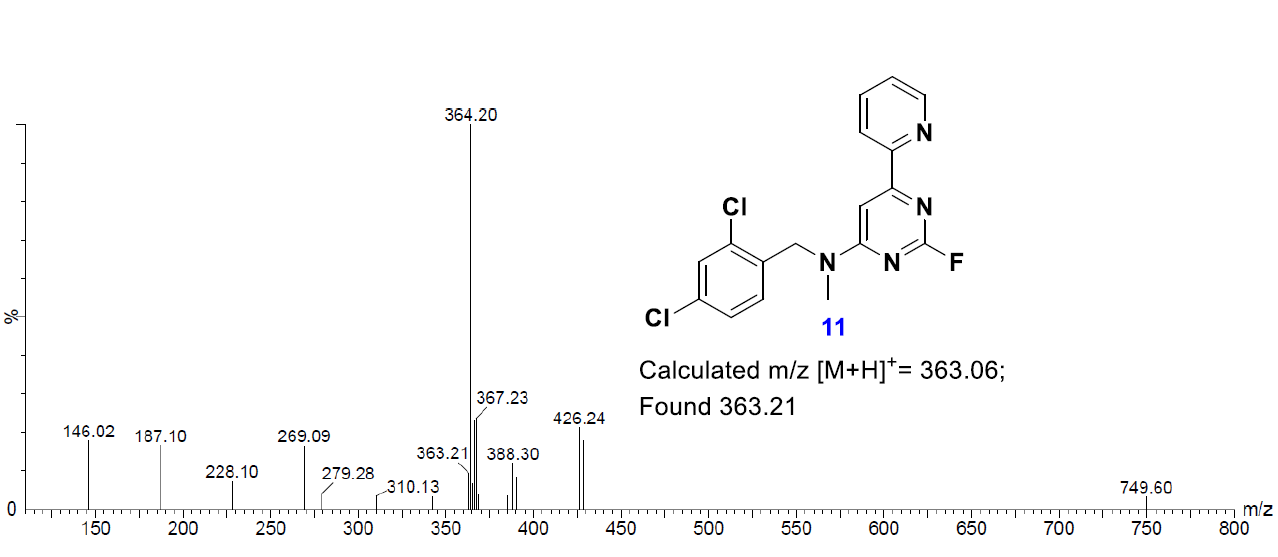

**Figure S14.** Mass spectrometry analysis of compound **11**.

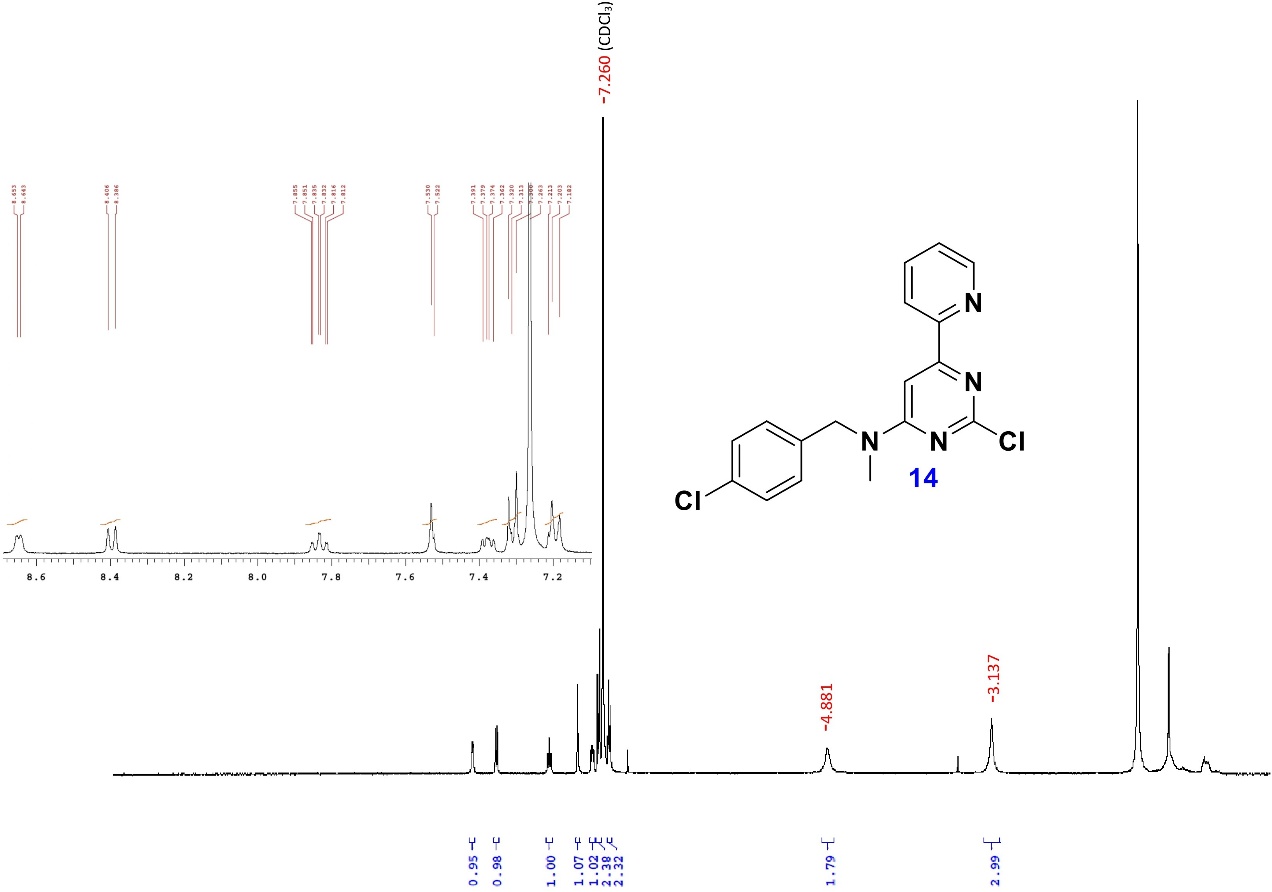

**Figure S15.** ^1^H NMR of compound **14**.

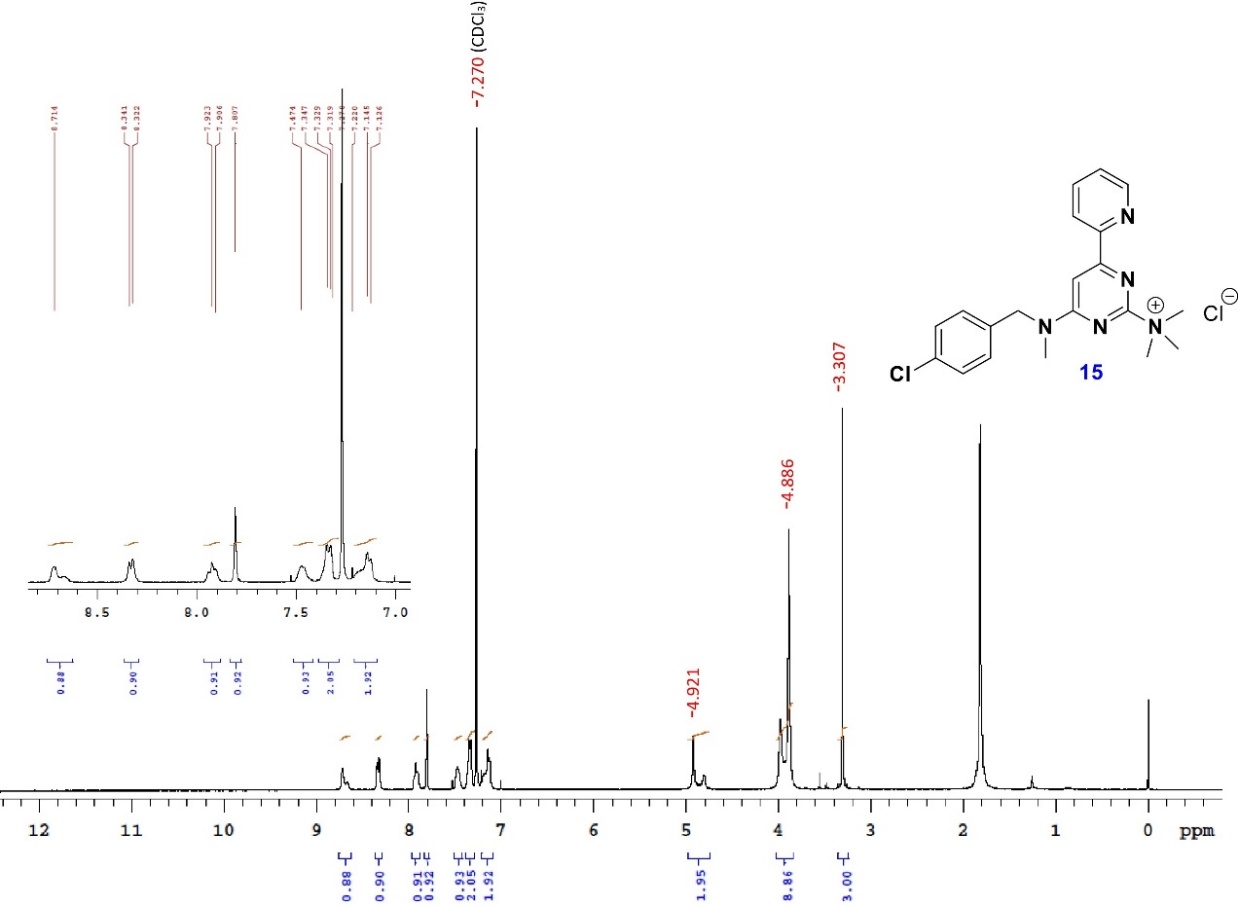

**Figure S16.** ^1^H NMR of compound **15**.

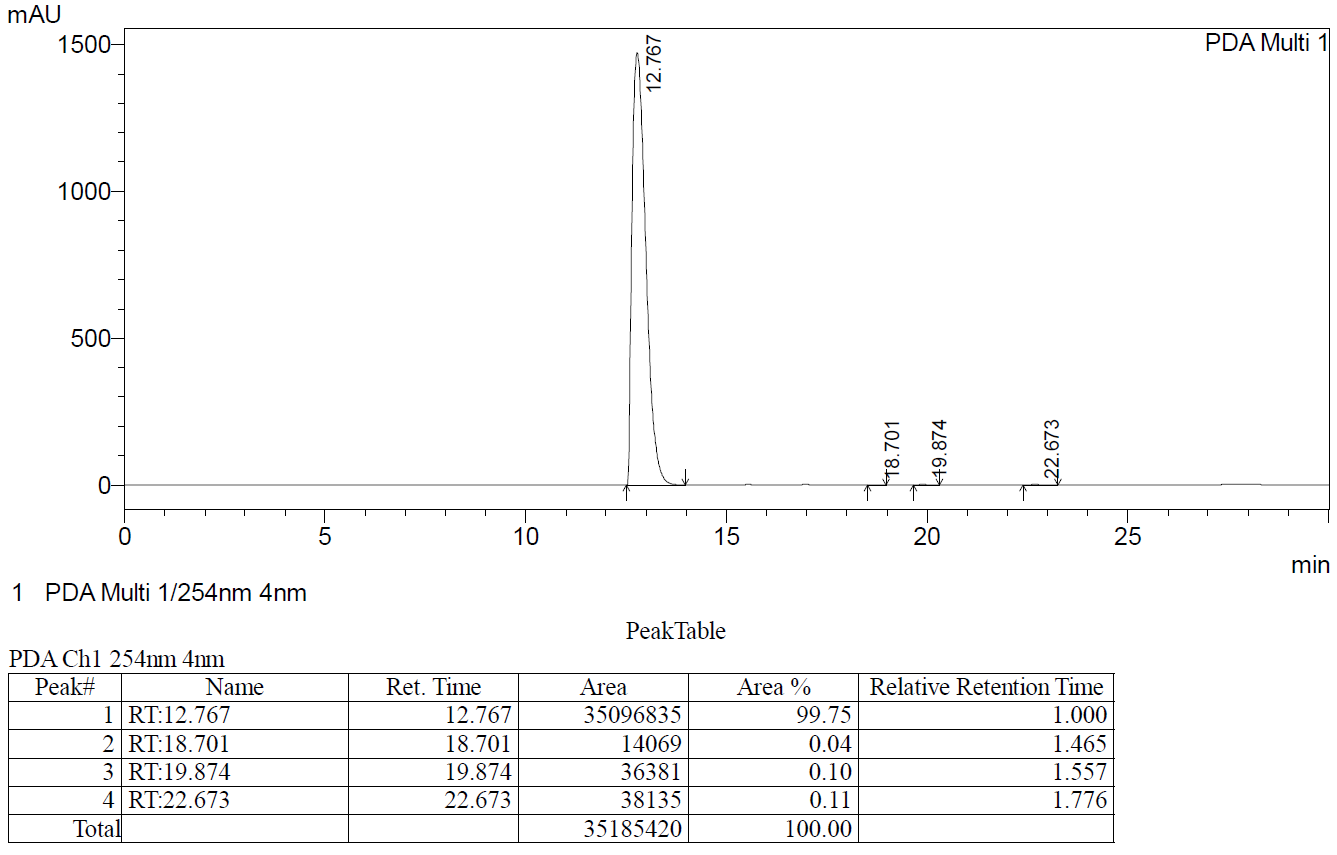

**Figure S17.** HPLC data of compound **15** (HPLC conditions: Column: Inertsil ODS-3V: 250 x 4.6 mm, 5 µm; mobile phase: 50% acetonitrile in water; flow rate: 1 mL/min).

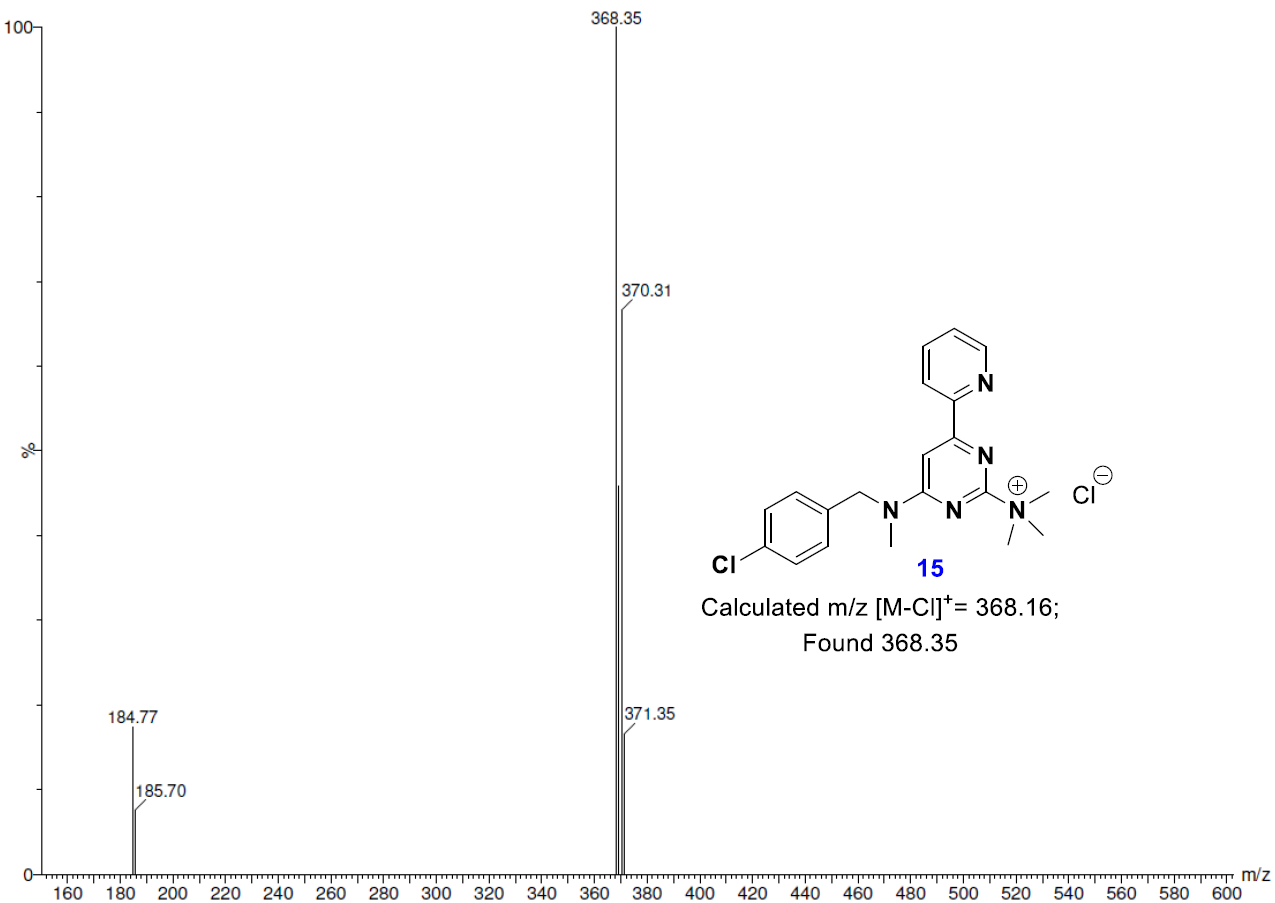

**Figure S18.** Mass spectrometry analysis of compound **15**.

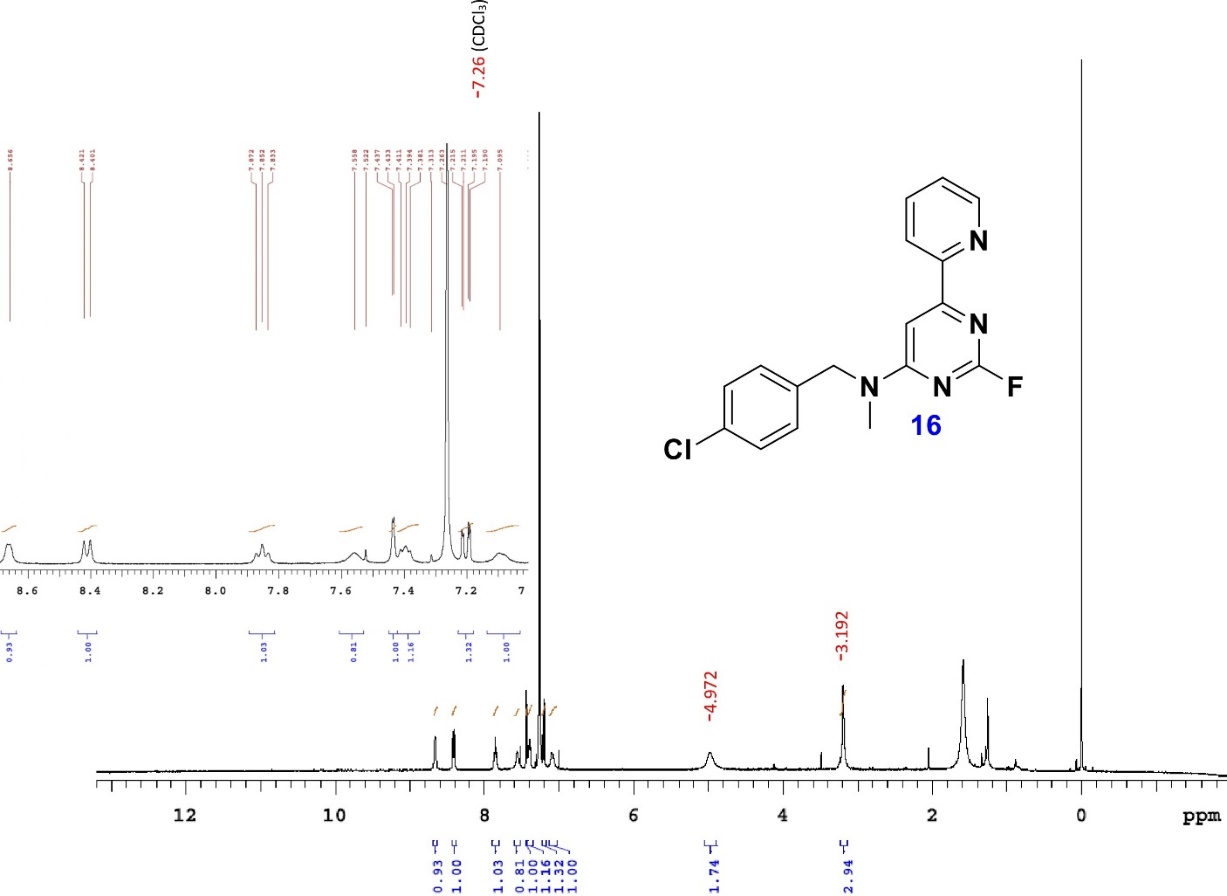

**Figure S19.** ^1^H NMR of compound **16**.

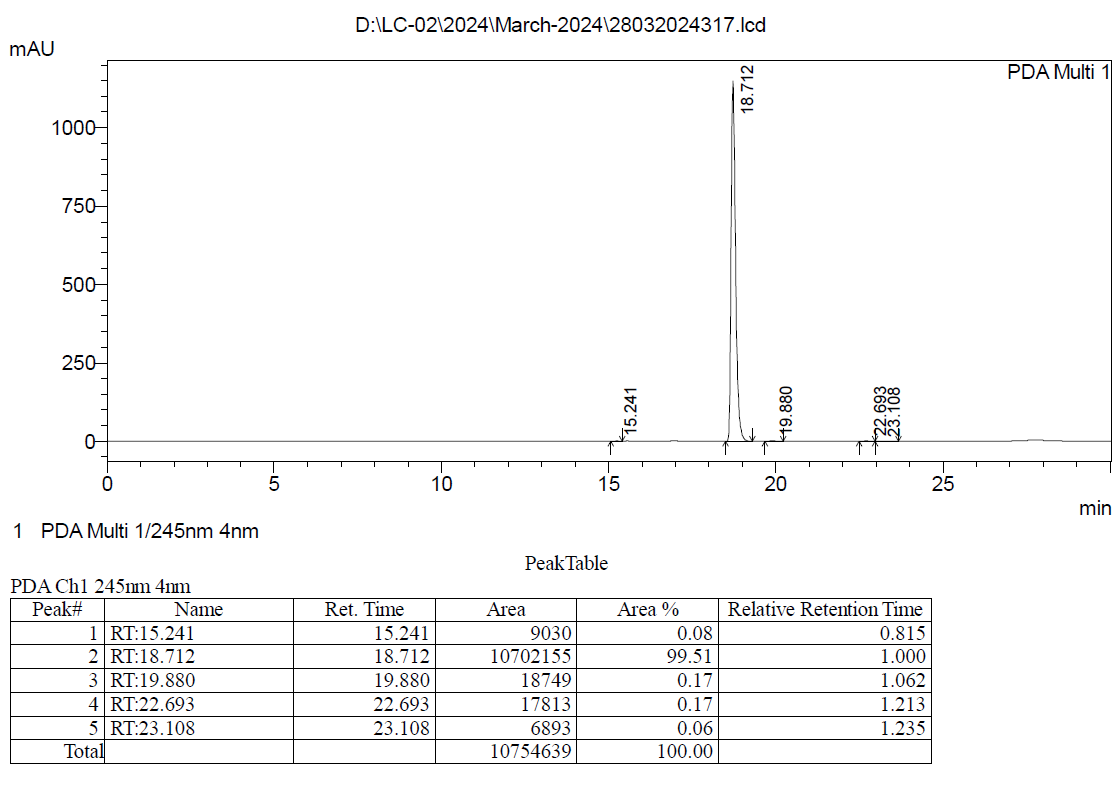

**Figure S20.** HPLC data of compound **16** (HPLC conditions: column: Inertsil ODS-3V: 250 x 4.6 mm, 5 µm; mobile phase: 50% acetonitrile in water; flow rate: 1 mL/min).

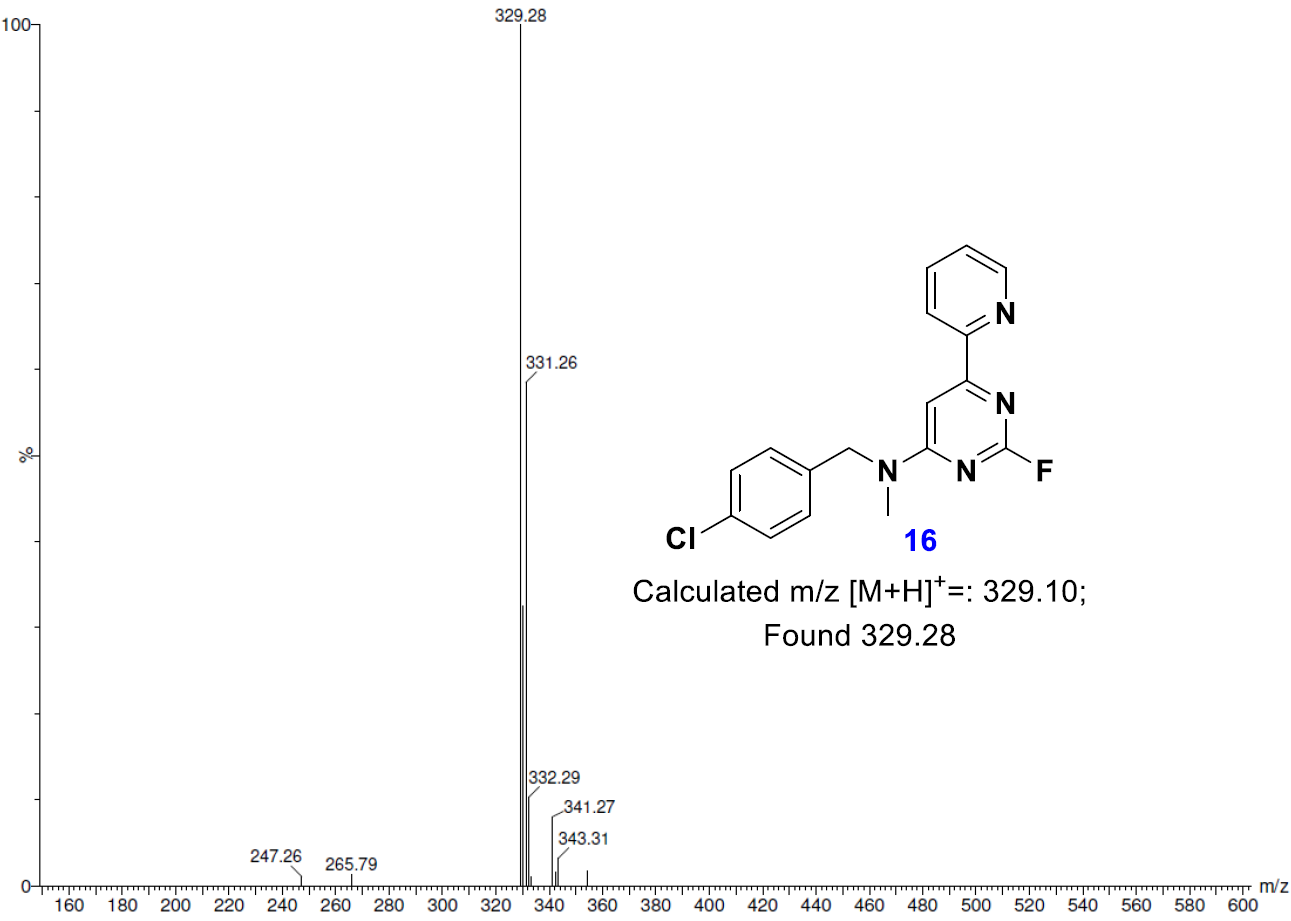

**Figure S21.** Mass spectrometry analysis of compound **16**.

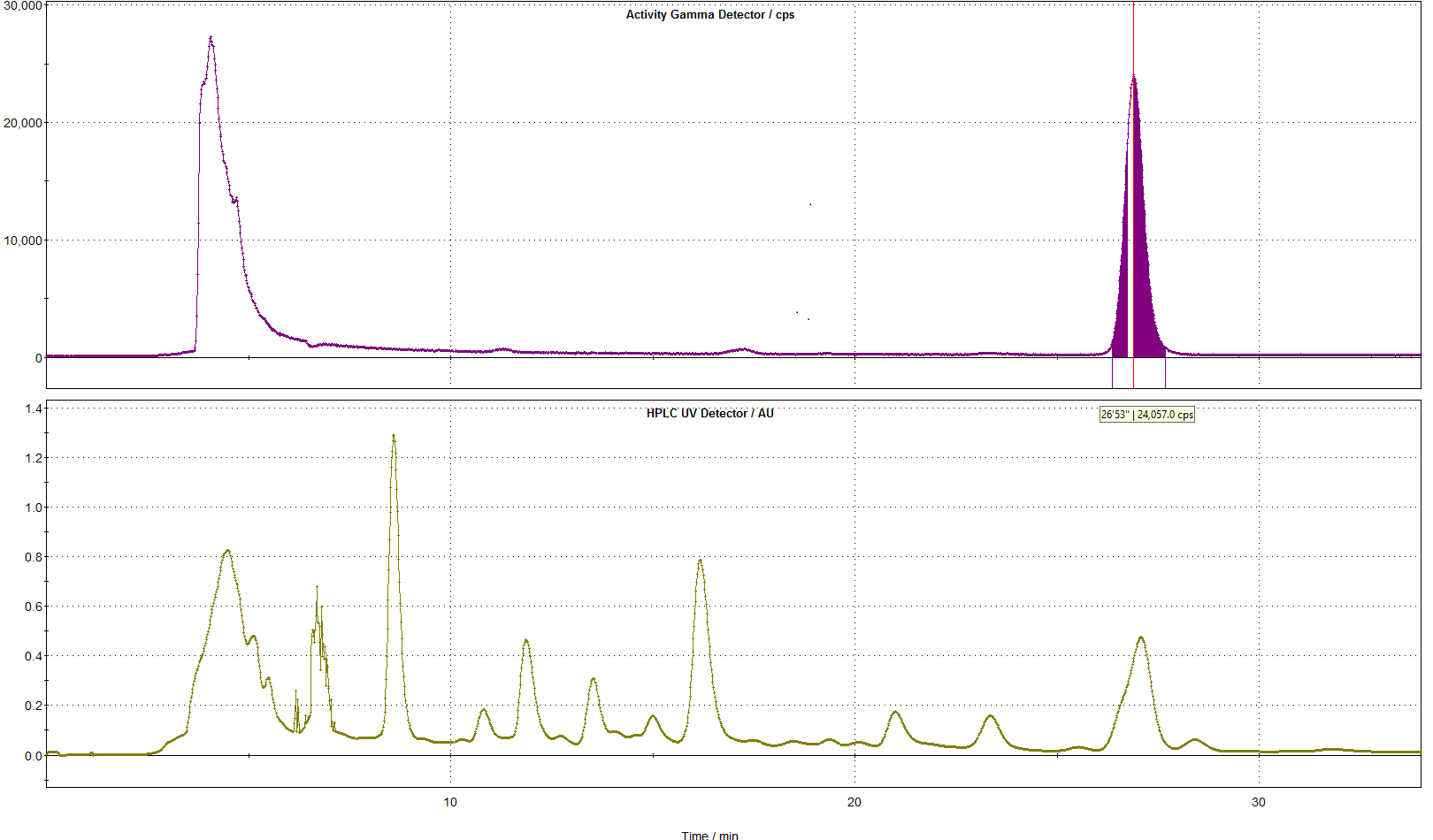

A. **Radioactivity channel**
B. **UV channel**

**Figure S22.** Representative semi-preparative HPLC chromatograms for [^18^F]GPR39-2.1 purification. (A) Radioactivity channel and (B) UV channel.

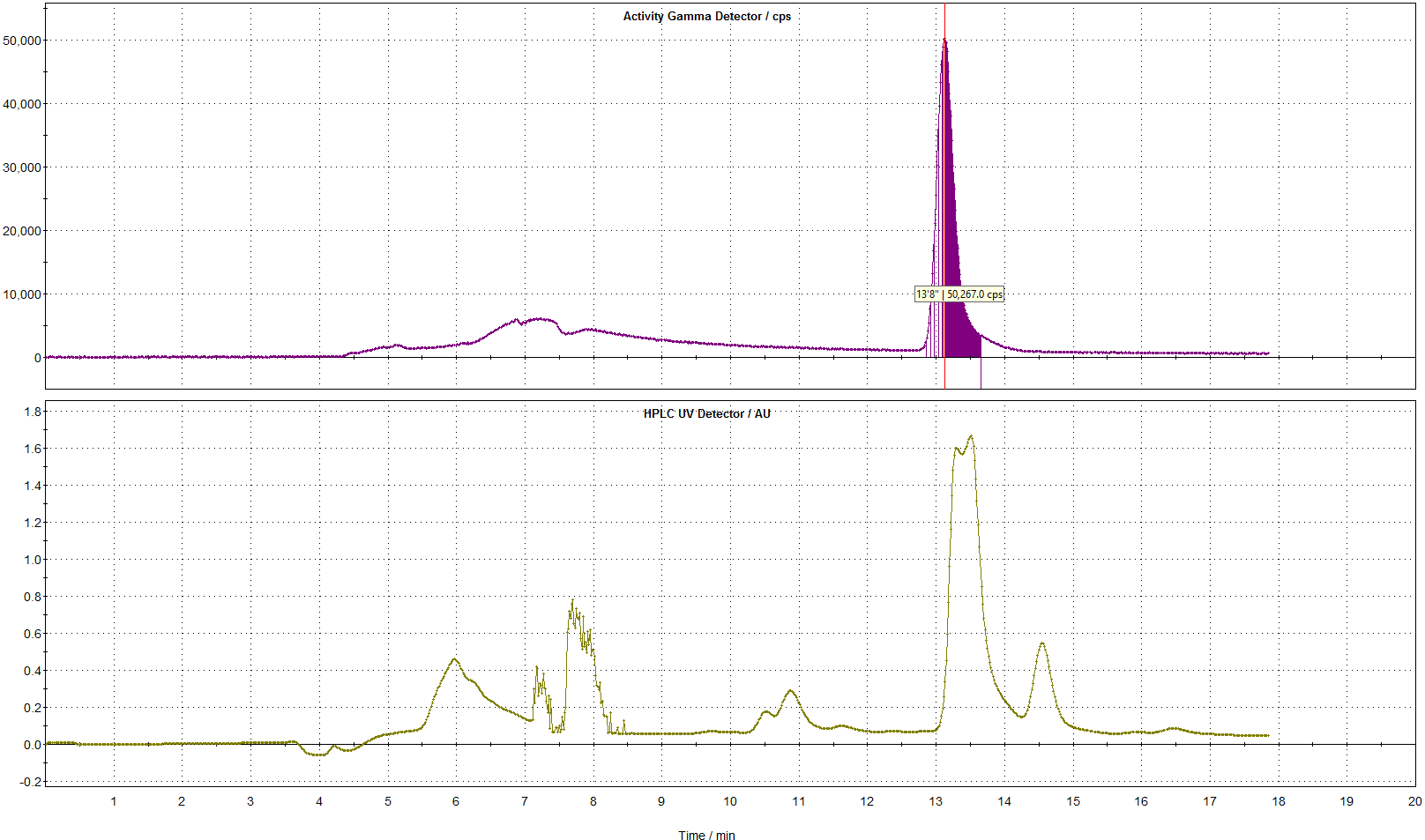

A. **Radioactivity channel**
B. **UV channel**

**Figure S23.** Representative semi-preparative HPLC chromatograms for [^18^F]GPR39-2.2 purification. (A) Radioactivity channel and (B) UV channel.

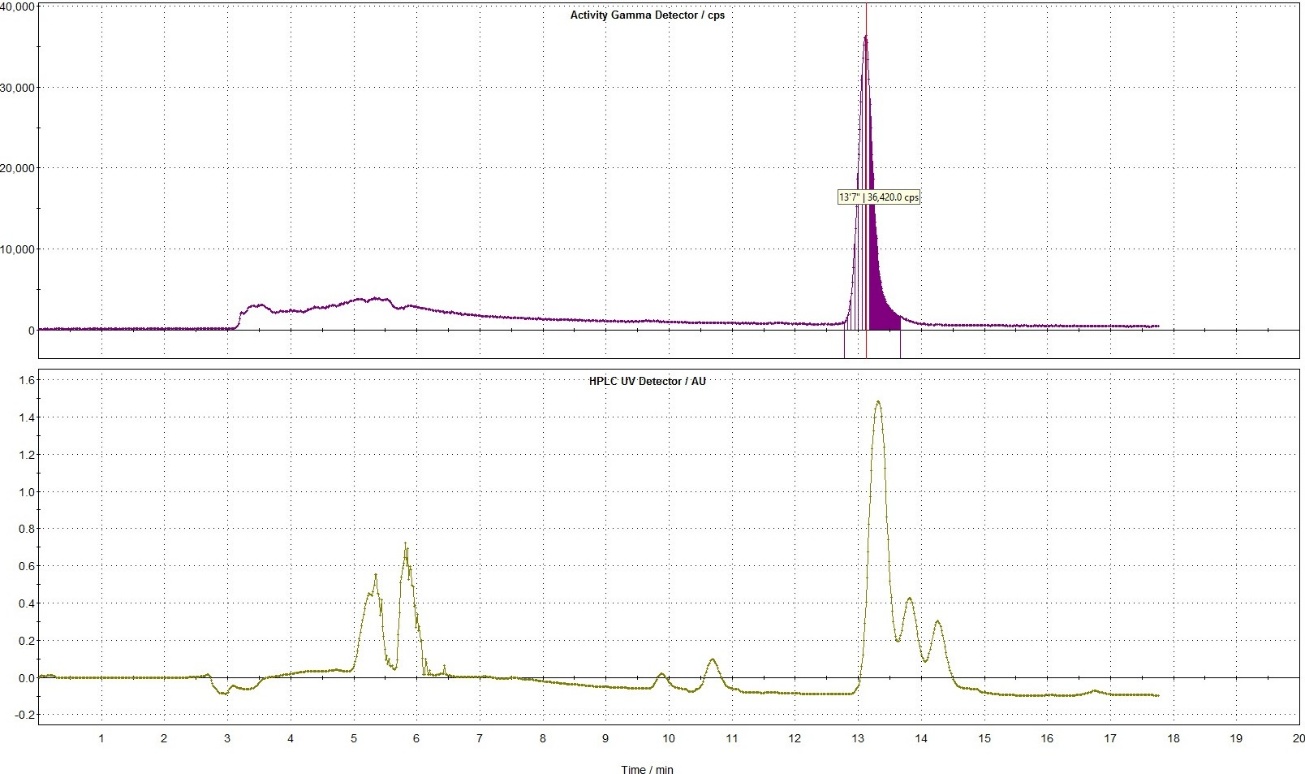

A. **Radioactivity channel**
B. **UV channel**

**Figure S24.** Representative semi-preparative HPLC chromatograms for [^18^F]GPR39-2.3 purification. (A) Radioactivity channel and (B) UV channel.

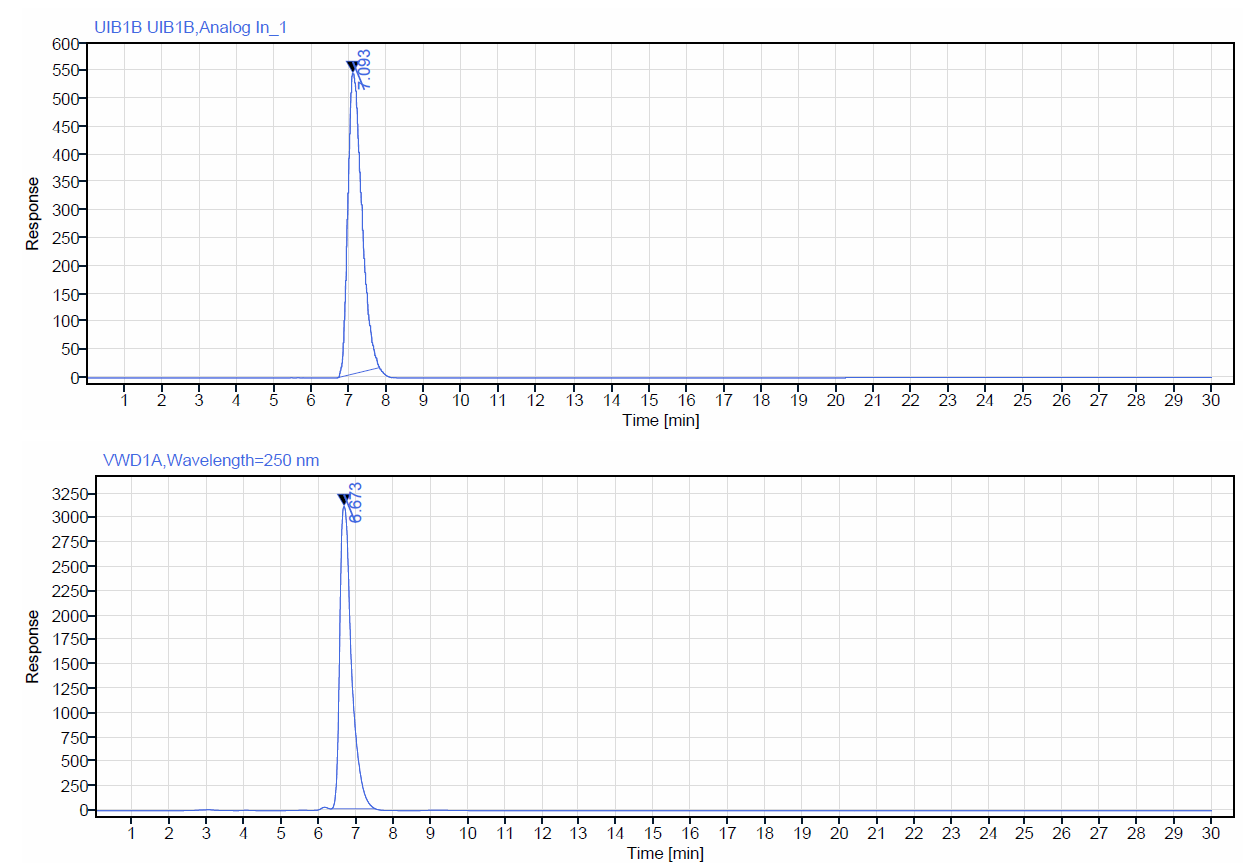

A. **Radioactivity channel**
B. **UV channel**

**Figure S25.** Representative analytical HPLC chromatograms of [^18^F]GPR39-2.1 and standard co-injection. (A) Radioactivity channel and (B) UV channel.

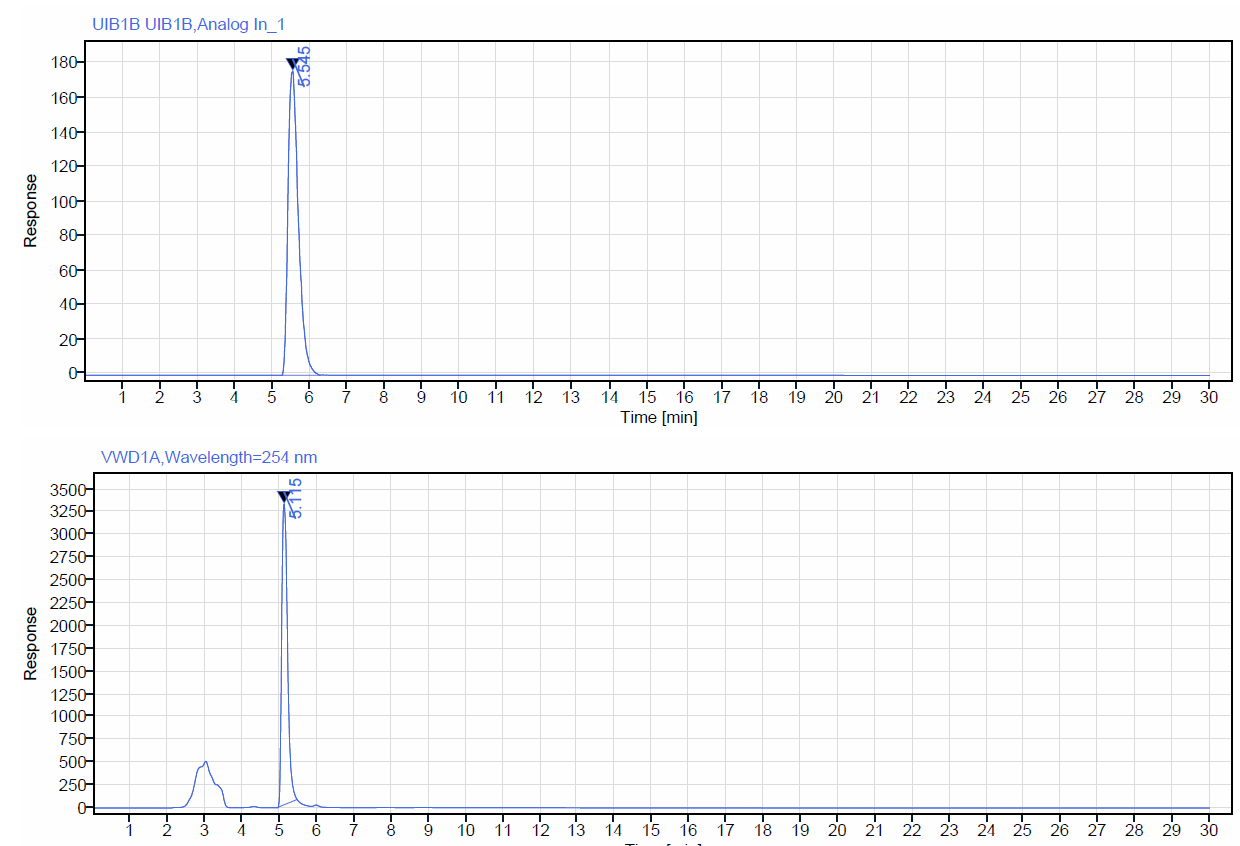

A. **Radioactivity channel**
B. **UV channel**

**Figure S26.** Representative analytical HPLC chromatograms of [^18^F]GPR39-2.2 and standard co-injection. (A) Radioactivity channel and (B) UV channel.

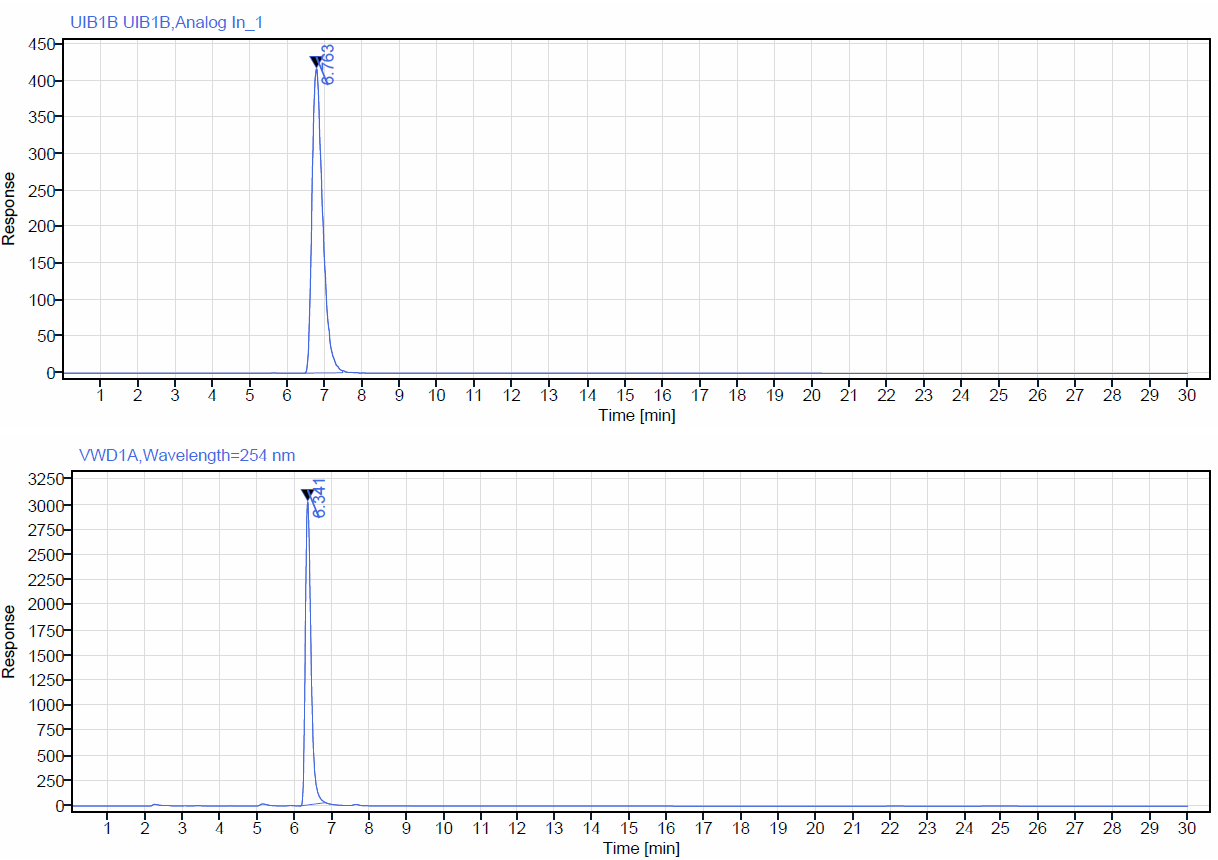

A. **Radioactivity channel**
B. **UV channel**

**Figure S27.** Representative analytical HPLC chromatograms of [^18^F]GPR39-2.3 and standard co-injection. (A) Radioactivity channel and (B) UV channel.

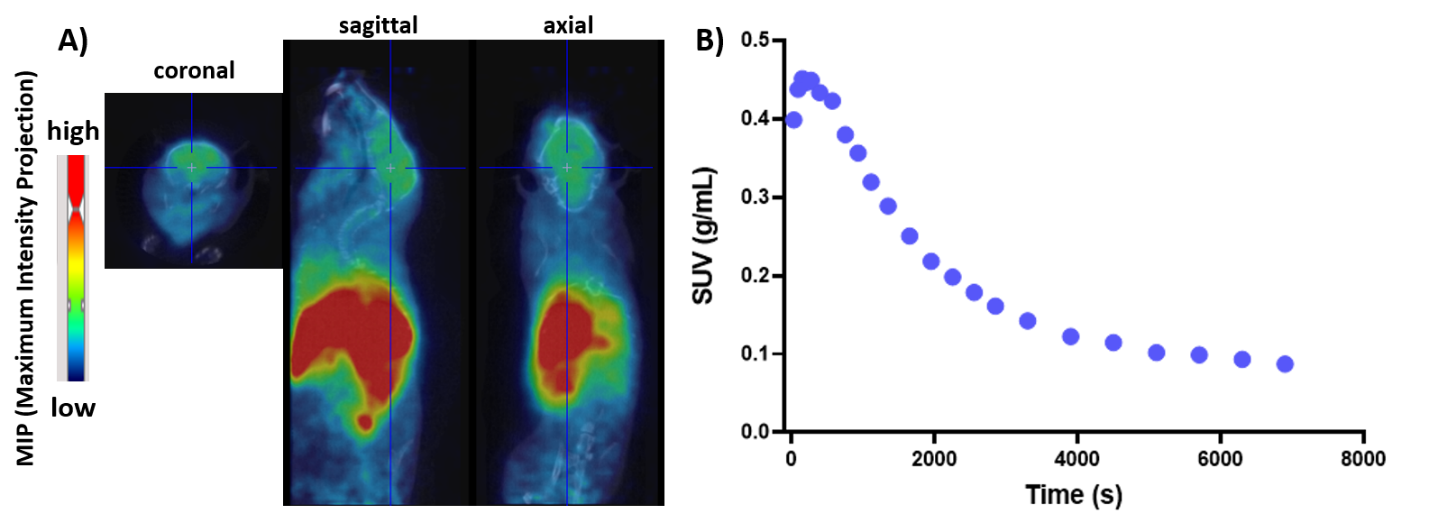

**Figure S28.** (A) Representative 2D images of [^18^F]GPR39-2.1 uptake in the whole body of BALB/c mice in coronal, sagittal, and axial views (B) Time activity curve of [^18^F]GPR39-2.1 in the whole brain over time (0-120 min) in BALB/c mice (n = 3).
